# TRPV4 antagonist efficacy in a post-hemorrhagic hydrocephalus pig model

**DOI:** 10.64898/2026.11.02.763674

**Authors:** R. Timothy Bentley, Scott C. Mitchell, Robyn R. McCain, Gregory T. Knipp, David Audu, Fatemeh O. Bidgoli, Alexandra Hochstetler, David J. Behm, Abigail D. Cox, Bonnie L. Blazer-Yost

## Abstract

Hydrocephalus occurs across the age spectrum but is particularly prevalent in young and old individuals. One form of this disease is post-hemorrhagic hydrocephalus (PHH), which is a life-threatening sequela of intraventricular hemorrhage (IVH). There are no effective non-surgical interventions for the treatment of PHH or other forms of hydrocephalus. Progress has been limited by the lack of gyrencephalic animal models that exhibit high fidelity to human conditions. Here, we establish a pathophysiologically relevant novel swine model of IVH that recapitulates key features of severe PHH, including sustained ventriculomegaly, elevated intracranial pressure (ICP), neuroinflammation and white matter injury. Using this model, we test the therapeutic potential of pharmacologic inhibition of the transient receptor potential vanilloid 4 (TRPV4) non-specific cationic channel. The orally bioavailable TRPV4 antagonist GSK2789917 penetrates the blood-brain barrier, and prevents progression to PHH when administered 2 days after IVH. Specifically, GSK2789917 prevents the development of ventriculomegaly, suppresses early ICP elevations and prevents or reverses inflammasome activation. These findings establish a clinically relevant large-animal model of PHH and identify TRPV4 antagonism as a promising treatment for hydrocephalus.

## Introduction

Hydrocephalus arises from many causes including hemorrhage, traumatic brain injury, infection, and neurodevelopmental syndromes. This spectrum of diseases affects patients from premature infants to senior citizens.^1–4^ With estimated incidences of 0.1% in the pediatric population and 1.5% in the geriatric population, hydrocephalus approaches Downs Syndrome and Parkinsonism respectively. Symptoms include pain, cognitive changes, vision, sleep and gait abnormalities. Hydrocephalus is often fatal if left untreated. Surgical diversion of excess cerebrospinal fluid (CSF) is the current treatment. However, the shunt placements used for this purpose exhibit high failure rates, routinely necessitating iterative brain surgeries. There is a substantial unmet global need for nonsurgical hydrocephalus therapies.

A major cause of hydrocephalus across ages is IVH, wherein the presence of blood in the ventricles disrupts normal CSF resorption, flow and/or production, leading to progressive ventriculomegaly and elevated ICP. PHH is a common sequala of IVH. Pharmacologic therapies that can be instituted immediately after IVH are a critical clinical need.

Our previous studies in a human choroid plexus epithelial (CPE) cell line and non-hemorrhagic rodent models showed that an antagonist of the electrolyte channel TRPV4 decreases CSF production and ameliorates hydrocephalus development, raising the question whether this therapy functions in a large animal preclinical model.^5,6^

Pharmaceutical development remains hampered by the lack of a physiologically relevant, large animal, gyrencephalic model encompassing high fidelity to human hydrocephalus including progressive ventriculomegaly, increased ICP and brain inflammation. To address this need, we modified a swine PHH model^7^, free from caustic chemicals, producing a model of sustained hydrocephalus exhibiting key hallmarks of the human condition.

In this high relevant model we tested an orally bioavailable TRPV4 antagonist and found it to be effective in mitigating PHH. The previously demonstrated human safety profile of this class of compounds^8–12^ makes them ideal for translation into human patients on the IVH-to-PHH continuum. Based on preclinical studies^5,6^, and the parallels between various forms of hydrocephalus including their responsiveness to surgical CSF diversion, these compounds could also be effective in hydrocephalus across multiple etiologies.

## RESULTS

### Development of a swine model of PHH

Yorkshire/Landrace cross barrows were selected to have a bodyweight of 25.0 – 31.6 kg, at the start of a 7-day acclimation period prior to each study. On Day 1 of a 31-day study (Fig. 1), IVH was induced by injecting a 12 ml solution (10 ml of autologous blood and 2 ml of 700 µm diameter embolism beads), into the right lateral ventricle (LV) in 21 pigs.

**Figure 1.**
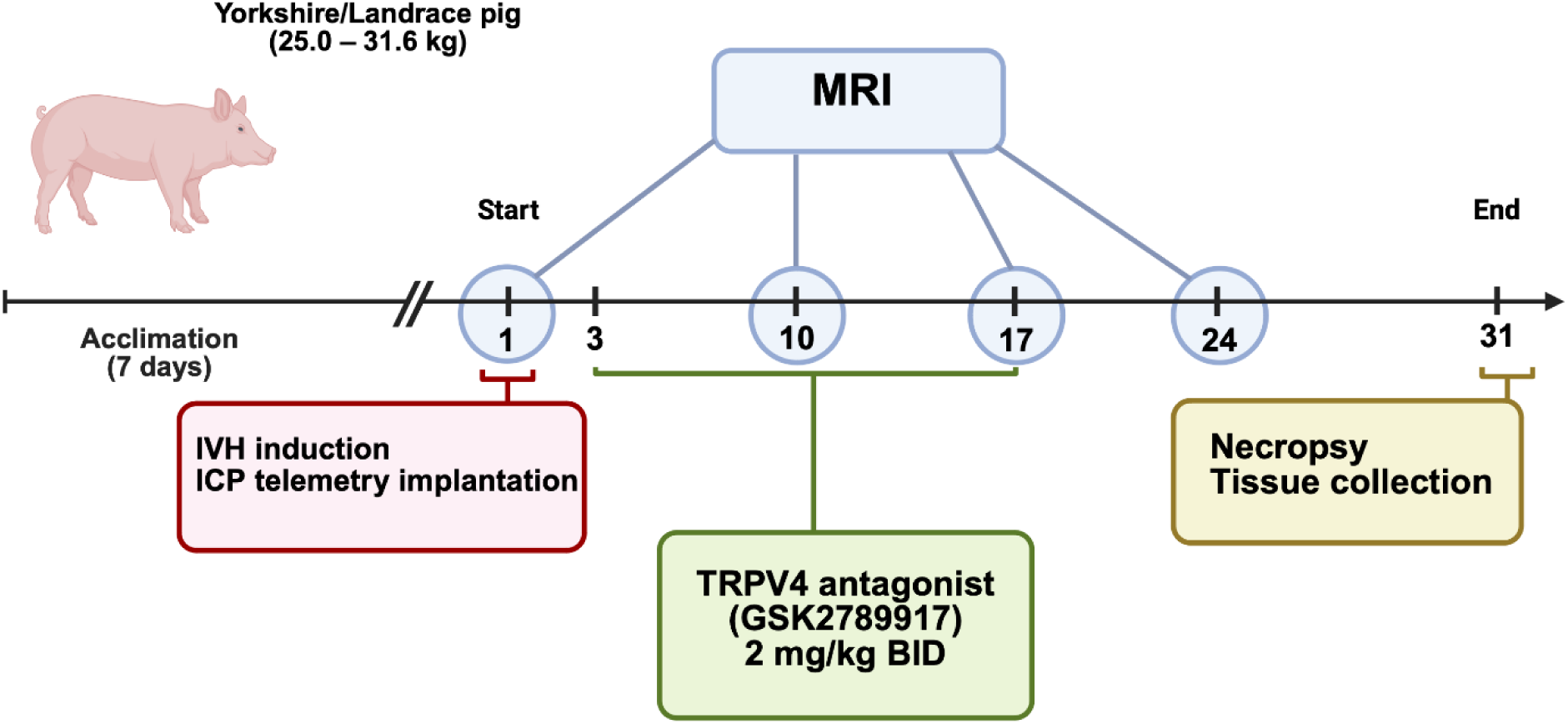
Experimental timeline of the swine IVH model with TRPV4 antagonist treatment. On Day 1, blood (10 mL) was drawn from the jugular vein and injected with embolism beads (2 mL) into the lateral ventricle. On Days 3 – 17, PHH-Treatment pigs (n = 6) received GSK2789917, but PHH-Control (n = 7) pigs did not. Four Sham-operated pigs received no blood injection, only an ICP probe placement. MRI was performed before IVH induction and at Days 10±1, 17±1 and 24±2. Abbreviations: ICP, intracranial pressure; IVH, intraventricular hemorrhage.

Ventricular size was measured using magnetic resonance imaging (MRI) on Days 1 (pre-IVH), 10, 17 and 24. ICP was measured throughout the study using implantable monitors placed immediately prior to IVH. A CT within 24 hr of surgery and the Day 10 MRI were evaluated for correct, intraventricular injection of the blood-beads mixture (Extended Data Fig. 1). 4 pigs for whom the LV was not accurately injected and 4 pigs that did not survive until 24 hours post-operatively were excluded. The 13 remaining pigs were allocated to the PHH-Control group (n = 7) or the PHH-Treatment group (n = 6). The PHH-Treatment group received an oral TRPV4 antagonist (GSK2789917) on Days 3 – 17. After sacrifice on Day 31, brains were harvested for histology. An additional 4 Sham-operated pigs (without blood–beads injection) provided normal values for ICP, ventricular size and pathology.

For the first three days following surgery, pigs were given a semi-quantitative neurological score based on examination by a board-certified veterinary neurologist. The morning after surgery, there was considerable variation in the severity of PHH pigs (Table 1). Two days later, all PHH pigs had 1 – 3 neurological deficits. On Day 22, one pig allocated to the PHH-Control group deteriorated and developed seizures. This subject was euthanized, without undergoing the final MRI.

**Table 1.** Neurological progression of pigs following surgery. Values represent the median number of neurological deficits detected (range).

|  | Morning after surgery<br>(Day 2) | Morning of Day 3<br>*Begin GSK*<br>*(Treatment group)* | Morning of Day 4 |
|---|---|---|---|
| Sham pigs<br>(n = 4) | 0 (0 – 1) | 0 (0 – 0) | 0 (0 – 0) |
| PHH-Control pigs<br>(n = 7) | 3 (1 – 7) | 3 (1 – 4) | 2 (1 – 3) |
| PHH-Treatment pigs<br>(n = 6) | 3 (2 – 4) | 3 (2 – 3) | 2 (2 – 3) |

Each of the following was recorded as a single deficit: reduced / delirious mentation; circling to the right; ataxic / paretic gait; left-sided menace deficit; unilateral proprioceptive deficits. Each of the following was recorded as two deficits: bilateral menace deficits; bilateral proprioceptive deficits.

### Brain Permeability and Pharmacokinetics

*In vitro* permeability analyses were used to determine whether the TRPV4 antagonist GSK2789917 permeates the blood brain barrier (BBB) and enters the neuronal chamber of the neurovascular unit (NVU). Using our established *in vitro* BBB and NVU models^13,14^, we conducted bidirectional permeability studies to ascertain whether GSK2789917 reaches brain parenchyma. The results (Table 2) revealed that GSK2789917 has a moderate rate of BBB permeation (the marker compound, acetaminophen, exhibits rapid permeation). Predicted steady-state concentration (C_ss_) is based on the influx and efflux permeabilities, and the efflux ratio and is used as a metric to assess brain accumulation.^15^ These *in vitro* results indicate that GSK2789917 penetrates the parenchyma of an unperturbed brain and achieve moderate therapeutic exposure.

**Table 2.** The *in vitro* BBB and NVU apparent permeability rates and the relative accumulation potential or predicted exposure (C_ss_) in brain parenchyma.

| **Compound and Model** | **Apparent Permeability (x10<sup>-6</sup> cm/sec)** |  | **Efflux Ratio (Bl→Ap/Ap→Bl)** | **Predicted Exposure Css** | **Permeability Rate** |
|  | Ap→Bl | Bl→Ap |  |  |  |
|---|---|---|---|---|---|
| GSK2789917 BBB | 2.59 ± 0.26 | 1.81 ± 0.44 | 0.70 | Medium to Low | Moderate |
| GSK2789917 NVU | 2.41 ± 0.55 | 1.48 ± 0.53 | 0.61 | Medium to Low | Moderate |
| Acetaminophen BBB | 13.1 ± 1.40 | 11.3 ± 1.18 | 0.86 | Medium | Fast |

Abbreviations: Ap , Apical; Bl, basolateral; C<sub>ss</sub>, steady-state concentration.

Next, we performed studies at doses of 2 and 4 mg/kg to evaluate single dose pharmacokinetic parameters in non-hydrocephalic pigs (8 samples, 0.25 – 12 hr after dosing). GSK2789917 does not appear to have a linear PK between single doses of 2 and 4 mg/kg (Table 3). The 4 mg/kg dose did not provide a true doubling of the C_max_ and AUC_0—Last_, although the T_max_ was much longer. A dose of 2 mg/kg, every 12 hours, was selected because it would provide sufficient blood levels after 24 hours to stay above the predicted EC_100_, and the T_max_ is faster. Accumulation will occur with multiple dosing, thus the concentration should plateau well above the EC_100_. Finally, single dosing appeared to provide levels that would elicit a predicted efficacious response to exposure for multiple doses.

**Table 3.** The calculated pharmacokinetic parameters using a noncompartmental model for single dosing of 2 and 4 mg/kg of GSK2789917 (2 male non-hydrocephalic pigs for each dose).

| **Dose** | **Parameter** | **Value (n = 2)** |
|---|---|---|
| 2 mg/kg | C<sub>MAX</sub> | 909 ng/mL |
|  | T<sub>MAX</sub> | 1.5 hr |
|  | T<sub>1/2</sub> | 3.3 hr |
|  | MRT<sub>0- t</sub> | 6.4 hr |
|  | AUC<sub>0-t</sub><sup>*</sup> | 3315.1 ng*hr/mL |
| 4 mg/kg | C<sub>MAX</sub> | 1332.9 ng/mL |
|  | T<sub>MAX</sub> | 8 hr |
|  | T<sub>1/2</sub> | 15.1 hr |
|  | MRT<sub>0- t</sub> | 11.2 hr |
|  | AUC<sub>0-t</sub><sup>*</sup> | 5038.9 ng*hr/mL |

Multiple dose pharmacokinetic parameters were also assessed in normal pigs. 9 samples per day, 0.25 – 24 hr after dosing, were drawn on 3 days (Table 4), with 3 samples per day on the other days. The noncompartmental PK parameters for 2 mg/kg BID dosing over 8 days revealed accumulation over time, with a potential C_ss_ occurring on day 5.

**Table 4.** The calculated pharmacokinetic parameters using a noncompartmental model for dosing 2 mg/kg BID of GSK2789917, in 4 male non-hydrocephalic pigs for the first 12 hours on days 1, 5 and 8.

| **Dose** | **Parameter** | **Day 1** | **Day 5** | **Day 8** |
|---|---|---|---|---|
| 2 mg/kg | C<sub>MAX</sub> (ng/mL) | 834 ± 127 | 1237 ± 69 | 1081 ± 275 |
|  | T<sub>MAX</sub> (hr) | 1.38 ± 0.25 | 2.3 ± 1.4 | 1.5 ± 0.0 |
|  | T<sub>1/2</sub> (hr) | 3.91 ± 0.64 | 5.4 ± 0.7 | 6.9 ± 1.6 |
|  | MRT<sub>0- t</sub> (hr) | 4.30 ± 0.8 | 5.1 ± 0.3 | 4.8 ± 0.7 |
|  | AUC<sub>0-t</sub> (ng /hr*mL) | 4836 ± 803 | 11128 ± 69 | 7527 ± 1768 |
|  | Kel (1/hr) | 0.18 ± 0.03 | 0.13 ± 0.02 | 0.11 ± 0.03 |

To determine whether hydrocephalus induces changes in the PK parameters, plasma was collected starting on Day 3 (first day of dosing), from 6 PHH-Treatment pigs (10 samples, 0.25 – 24 hr after morning dose). Two were omitted from PK analysis, due to seizures and seizure therapy possibly interfering with GSK2789917 bioavailability. The parameters of the remaining four PHH pigs (Table 5) were similar to those on day 5 of the multiple dosing study, suggesting that IVH did not alter pharmacokinetics.

**Table 5.** The calculated pharmacokinetic parameters using a noncompartmental model for dosing 2 mg/kg of GSK2789917 BID on Day 3, in 4 male PHH pigs (first 24 hours after the morning dose) appeared similar to those on day 5 in naïve pigs.

| **Dose** | **Parameter** | **Value (n=4)** |
|---|---|---|
| 2 mg/kg | C<sub>MAX</sub> | 1409 ± 135 ng/ml |
|  | T<sub>MAX</sub> | 2.4 ± 1.1 hr |
|  | T<sub>1/2</sub> | 8.7 ± 2.1 hr |
|  | MRT<sub>0-t</sub> | 5.7 ± 0.6 hr |
|  | AUC<sub>0-t</sub> | 11443.3 ng/hr*mL |

### GSK2789917 inhibits ventriculomegaly

MRI was performed immediately before surgery (Day 1), and on Days 10, 17 and 24 to coincide with 1-week and 2-weeks of medication, and 1-week after discontinuing medication. Pre-operatively, there was no significant difference between any cohorts in Total Ventricular Volume (Fig. 2a & 2d) or the volumes of individual ventricles (Fig. 2c) (Extended Data Table 1).

**Figure 2.**
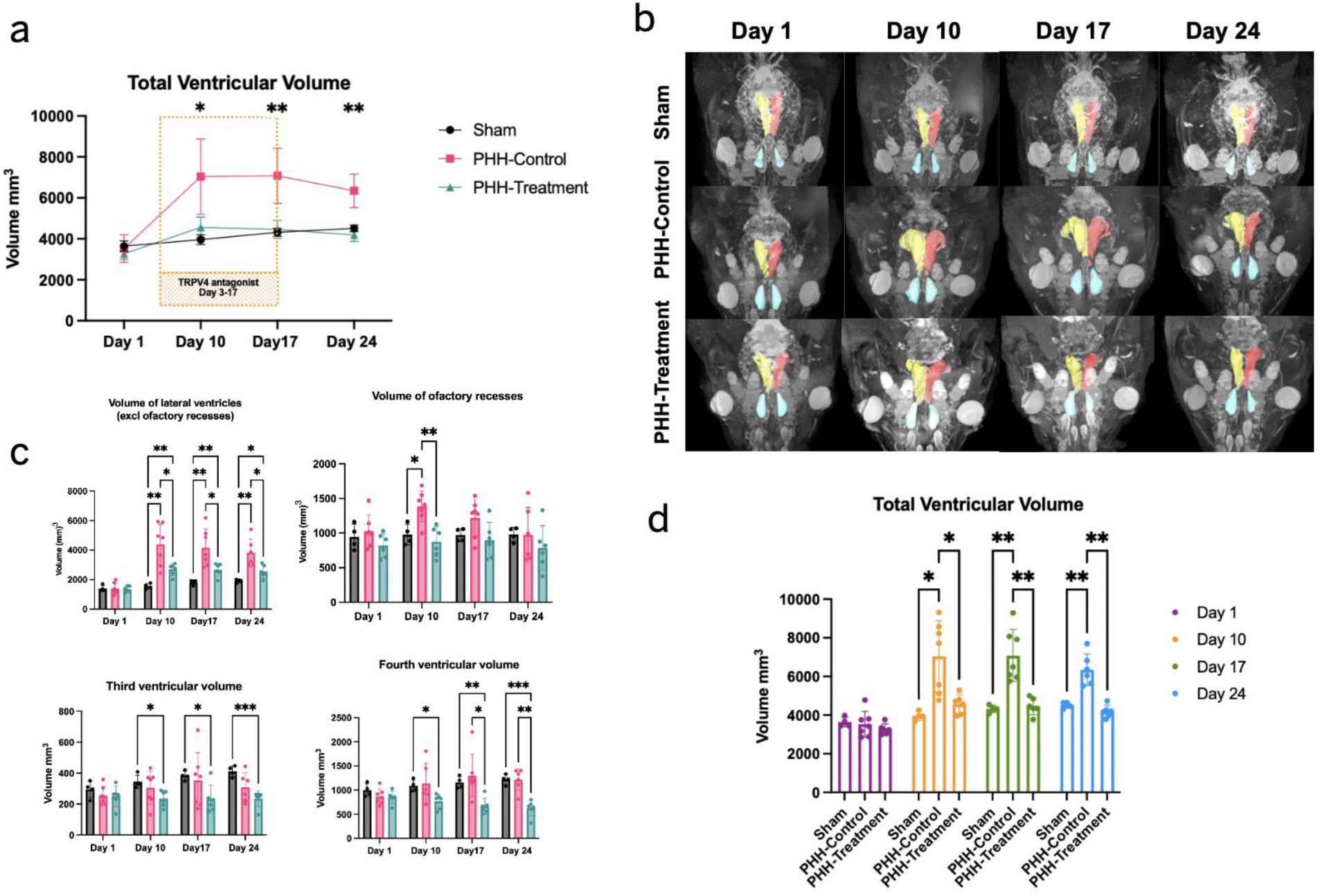
Magnetic resonance imaging (MRI) of swine post-hemorrhagic hydrocephalus (PHH), with and without TRPV4 antagonist treatment. PHH-Control and PHH-Treatment animals underwent surgery to induce PHH on Day 1. MRIs were performed pre-operatively, and at Days 10, 17 and 24. PHH-Treatment animals received the TRPV4 antagonist GSK2789917 on Days 3 – 17. (a) PHH-Treatment animals had significantly smaller Total Ventricular Volume than PHH-Control, on all post-operative MRIs. (b) Representative MRI examples of each cohort. The right LV is depicted in yellow and the left LV is depicted in red. The olfactory recesses of the lateral ventricles, which are large in pigs, are depicted in blue. (c) Volumes of the lateral ventricles, the olfactory recesses, the 3^rd^ ventricle and the 4^th^ ventricle revealed multiple significant differences between the Sham, PHH-Control and PHH-Treatment groups. These volumes were summed to create the Total Ventricular volume in Panel (a). (d) Total Ventricular Volume. There was no significant difference between the Sham and PHH-Treatment groups at any time. The PHH-Control group had significantly larger ventricles at all post-operative timepoints. Mean (box) and standard deviation (whiskers). *p < 0.05, **p < 0.01, ***p < 0.001.

In Sham-operated subjects, ventricular volumes increased slightly with growth (Fig. 2a). In both PHH-Treated and PHH-Control, blood was injected into the right LV, and the right ventricle was larger and contained more blood on Day 10 (Supplementary Fig. 1). Blood also entered the left LV. Negligible blood was observed in the aqueduct, 3^rd^ and 4^th^ ventricles, which were not enlarged by PHH induction (PHH-Control compared to Sham, Fig. 2c).

In the PHH-Control group, there was a considerable increase in ventricular volumes between the pre-operative MRI and the Day 10 MRI (Fig. 2a-2d). The lateral ventricles almost doubled in size. The significant enlargement of the lateral ventricles and Total Ventricular Volume was sustained on Days 17 and 24, compared to Sham.

The PHH-Treated group received GSK2789917 on Days 3 – 17. Compared to the PHH-Control group, they had statistically significantly smaller Total Ventricular Volumes post-operatively (Fig. 2a & 2d). Surprisingly, this significant difference persisted one week after discontinuing GSK2789917. Furthermore, there was no significant difference in the Total Ventricular Volume between PHH-Treated and Sham-operated at any timepoint (Fig. 2d) indicating that the TRPV4 antagonist was effective in preventing ventriculomegaly, with a sustained effect after discontinuing therapy.

In the PHH-Control group, the pig with the largest ventricles (pre- and post-operatively) was sacrificed on Day 22 due to neurological deterioration and seizures, prior to the final MRI. The decrease in the Control group mean Total Ventricular Volume from Day 17 to 24 should be interpreted in light of the removal of this individual.

### GSK2789917 alters PHH ICP trajectory

3 Sham, 6 PHH-Control, and 4 PHH-Treatment animals had epidural ICP readings that exhibited stereotyped waveforms for the duration of the study, including respiratory and cardiac cycles (Fig. 3), sufficient for analysis. In pigs where ICP was elevated over several days (Fig. 3b), this often coincided with a decrease in activity and fluctuation in temperature. In animals with enclosed calvaria, like pigs and humans, ICP measurements are generally considered normal from 5-20 mmHg, moderately elevated from 20-40 mmHg, and critically elevated >40 mmHg. One PHH-Control animal had critically elevated ICP for the first week. This pig was especially lethargic on Days 2 and 7, initially improved, then was euthanized at Day 22 due to sudden recrudescence.

**Figure 3:**
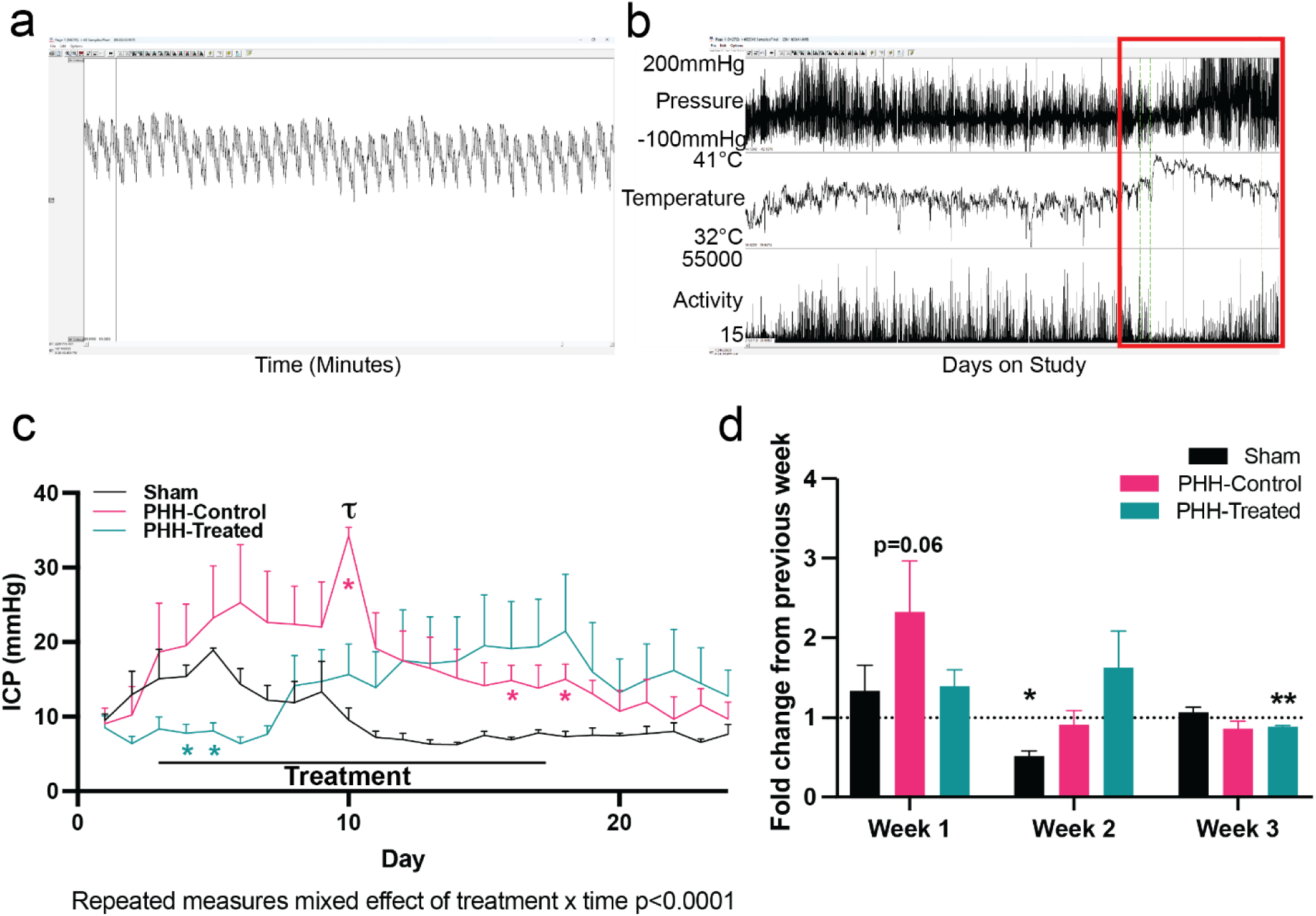
Continuous epidural intracranial pressure (ICP) monitoring in pigs. (a) Representative ICP waveform collected with an implanted epidural monitor. Note the slow oscillation from the respiratory cycle, and the superimposed more rapid oscillation consistent with the cardiac cycle. (b) Example of a case where ICP (top) was elevated over several days (red box), accompanied by changes in temperature (middle) and pig activity (lower). (c) Mean + standard error daily values for ICP. GSK2789917 was administered to the PHH-Treated group on Days 3 – 17. The 3 groups were statistically significantly different (repeated measures P < 0.0001). Asterisks indicate p<0.05 compared to sham. Tau indicates p<0.05 compared between PHH groups. (d) Changes to ICP presented as fold change from baseline (Days 1-2) on a week-by-week basis (Days 3-10 = “Week 1”, Days 11-17 = “Week 2”, Days 18-24 = “Week 3”). Asterisks indicate *p<0.05; **p<0.01 in one-sample Wilcoxon test compared to preceding time period.

In Figure 3c, data are presented as daily mean ICP for each group. A linear mixed-effects model revealed a significant interaction between treatment group and time (p < 0.0001), indicating that ICP changes differed significantly across groups during the study. It can be appreciated that GSK2789917 prevented the initial spike in ICP after IVH. In addition, at several discrete timepoints, post-hoc multiple t-tests with Holm’s Sitak correction observed a significant difference (p<0.05) between PHH-Control and Sham animals (pink asterisk), between PHH-Treatment and Sham animals (teal asterisk), and between PHH-Control and PHH-Treatment (tau symbol). When presented as fold change from baseline for each group (Fig. 3d), GSK2789917 significantly prevents early ICP elevations. Together, these findings demonstrate that treatment is associated with a lower initial ICP and significantly alters the trajectory of ICP over time, leading to divergence of treated and untreated PHH animals during and after treatment.

### GSK2789917 ameliorates PHH etiopathology

Structural and neuroinflammatory alterations in the choroid plexus (CP), ependyma and periventricular brain tissue of PHH pigs were significantly reduced by TRPV4 antagonist therapy. Evaluation of the inflammatory response to PHH occurred 31 days after the hemorrhagic insult and two weeks after the final dose of GSK2789917.

The structural impact of PHH and GSK2789917 treatment was evaluated via hematoxylin and eosin (H&E) staining (Fig. 4 & 5). Histological examination of Sham CP revealed a normal architecture characterized by thin fibrovascular cores and regular capillary networks (dotted black arrow) (Fig. 4a). Similarly, the Sham LV exhibited normal morphology with an intact, continuous ependymal lining (black arrow) (Fig. 5a). Untreated PHH-Control CP tissue exhibited marked morphological alterations, substantial CP stromal edema and severe vascular dilation (black arrow) (Fig. 4a). This group demonstrated marked ventriculomegaly, accompanied by severe stretching and disruption of the ependymal cell layer along the ventricular wall (dotted black arrow) (Fig. 5a, c). Treatment of PHH-induced pigs with GSK2789917 attenuated these pathological changes, resulting in structural CPE preservation more closely resembled the healthy architecture of the Sham controls (Fig. 4a), partially attenuated ependymal disruption and reduced ventricular size to untreated PHH-Controls (Fig. 5a, c).

**Figure 4.**
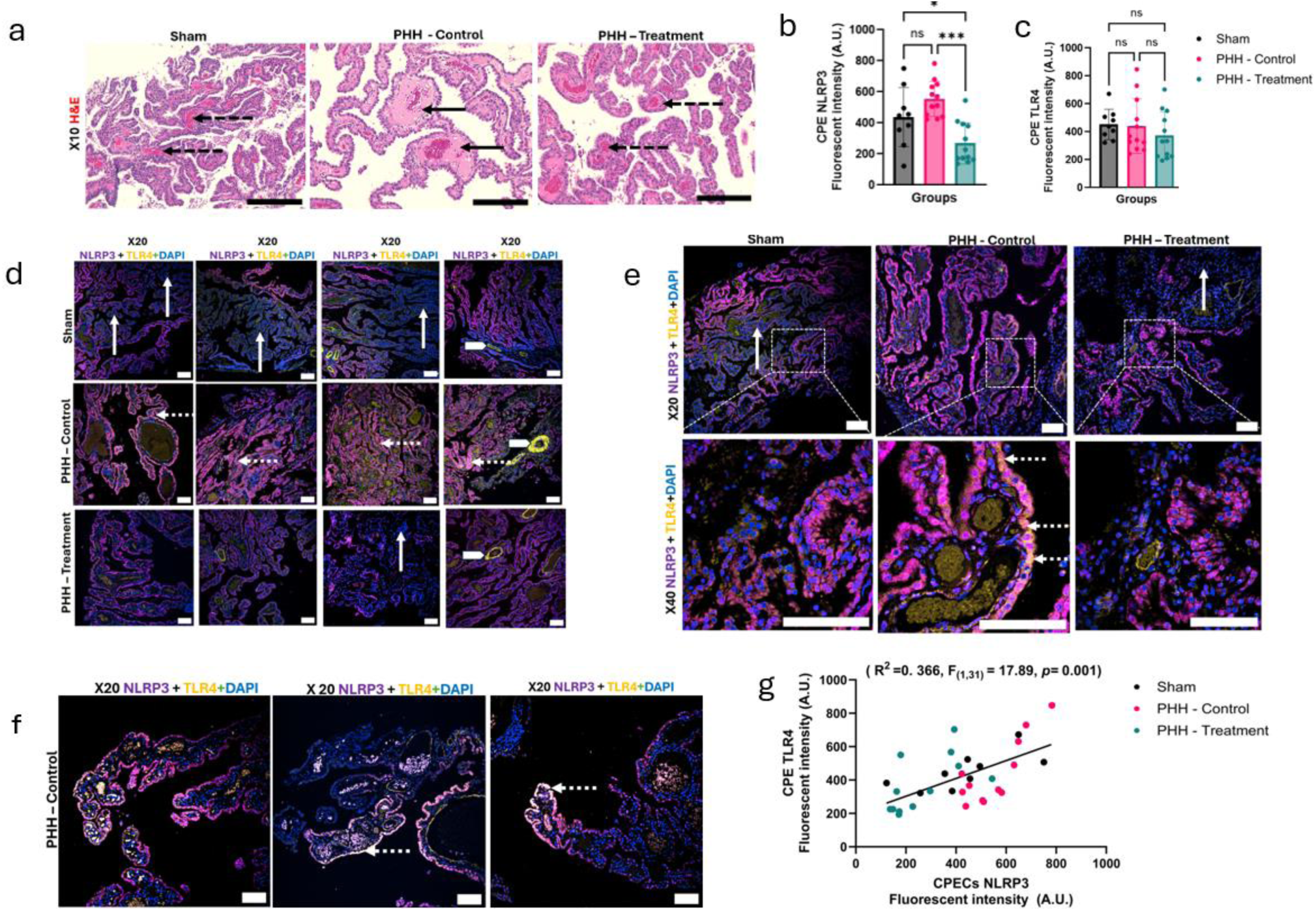
Histological and neuroinflammatory assessment of the choroid plexus (CP) (a) Representative hematoxylin and eosin (H&E) images of the choroid plexus (CP). Sham animals exhibit preserved CP architecture characterized by thin fibrovascular cores and organized capillary networks (dotted black arrow). PHH-Control CP demonstrates stromal edema and prominent vascular dilation (black arrow). Treatment with GSK2789917 attenuates these structural alterations, with partial restoration of normal CP morphology (dotted black arrow). Scale bar: 50 µm. (b) Quantification of NLRP3 fluorescence intensity (arbitrary units, A.U.) in the CP. (c) Quantification of TLR4 fluorescence intensity (A.U.) in the CP. (d–f) Representative confocal immunofluorescence images of CP sections from Sham, PHH-Control, and PHH-Treatment animals stained for NLRP3 (magenta), TLR4 (yellow), and DAPI (blue). White arrows indicate regions of low basal NLRP3 and TLR4 expression within the choroid plexus epithelium (CPE). White block arrows denote vascular TLR4 localization. Dotted white arrows highlight regions of NLRP3 and TLR4 colocalization. Scale bars: 50 µm (low magnification) and 20 µm (high magnification). (g) Linear regression analysis of CPE TLR4 versus NLRP3 fluorescence intensity across individual samples from all experimental groups: Sham, n = 3; PHH-Control, n = 4; and PHH-Treated, n = 4. 3 distinct aspects of the same tissue per animal were analyzed. Data = mean ± SEM. *p < 0.05, ** p < 0.01, *** p < 0.001. Abbreviations: CP, choroid plexus; CPE, choroid plexus epithelium; PHH, post-hemorrhagic hydrocephalus; NLRP3, NOD-like receptor family pyrin domain-containing 3; TLR4, Toll-like receptor 4; DAPI, 4′,6-diamidino-2-phenylindole.

**Figure 5.**
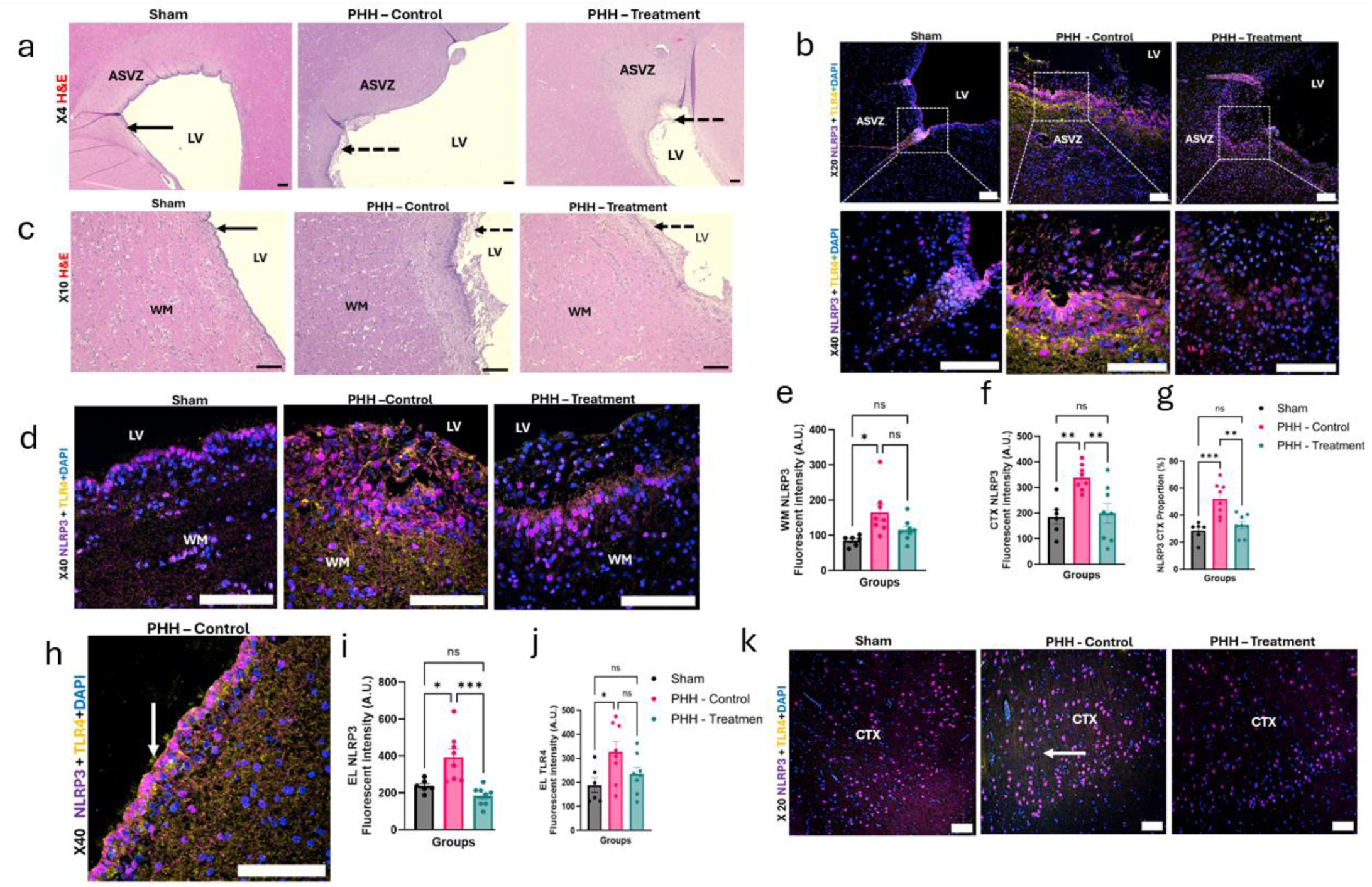
Structural and neuroinflammatory characterization of the ventricular system, white matter, and cortex. (a,c) H&E images of the lateral ventricle (LV), subventricular zone (ASVZ), and white matter (WM) in Sham, PHH-Control, and PHH-Treatment pigs. Sham shows an intact ependymal lining (black arrows), whereas PHH-Control and PHH-Treatment show ependymal disruption (dotted black arrows); ventricular (LV) dilation and WM degradation are most pronounced in PHH-Control. Scale bars: 200 µm (LV and ASVZ) and 100 µm (WM). (b,d) Confocal images of LV, anterior ASVZ, and periventricular WM, stained for NLRP3 (magenta), TLR4 (yellow), and DAPI (blue). Dashed boxes indicate higher-magnification LV and ASVZ panels below. Scale bars: 50 µm (upper LV/ASVZ panels), 20 µm (lower LV/ASVZ panels), and 20 µm (WM). (e,f) Quantification of NLRP3 fluorescence intensity (arbitrary units, A.U.) in WM and cortex (CTX). (g) Quantification of cortical NLRP3-positive cell proportion (% of CTX). (h) Confocal image of ependymal lining in PHH-Control tissue stained for NLRP3 (magenta) and TLR4 (yellow). Arrow indicates cilia with TLR4 localization. Scale bar: 20 µm. (i,j) Quantification of ependymal NLRP3 and TLR4 intensity (A.U.). (k) Confocal immunofluorescence images of cortex (CTX) stained for NLRP3 (magenta), TLR4 (yellow), and DAPI (blue). White Arrow indicates TLR4 localization. Scale bar: 50 µm. n = 3 pigs (Sham), n = 4 pigs (PHH), and n = 4 pigs (PHH-treated); 2 distinct aspects of the same tissue per animal were analyzed for the EL, WM, and CTX. Data = mean ± SEM. *p < 0.05, ** p < 0.01, *** p < 0.001. **Abbreviations:** ASVZ, anterior subventricular zone; CTX, cerebral cortex; DAPI, 4′,6-diamidino-2-phenylindole; EL, ependymal lining; LV, lateral ventricle; NLRP3, NOD-like receptor family pyrin domain-containing 3; TLR4, Toll-like receptor 4; WM, white matter.

The periventricular white matter (WM) in the Sham cohort displayed compact, organized parenchyma (Fig. 5c). In contrast, PHH-Control animals exhibited pronounced WM structural degradation.

Treatment with GSK2789917 improved the preservation of WM integrity (Fig. 5c). Within deeper brain parenchyma (Supplementary Fig. 2f), Sham cerebral cortex (CTX) exhibited preserved cytoarchitecture with uniform cellular organization. In contrast, untreated PHH-Control cortex demonstrated vacuolation (black arrow). Treatment with GSK2789917 attenuated cortical vacuolation and improved structural preservation, resulting in cytoarchitecture comparable to Sham controls.

To visualize the spatial distribution and cellular localization of the inflammatory response, we performed immunofluorescence staining for NOD-like receptor family pyrin domain-containing 3 (NLRP3) (magenta) and Toll-like receptor 4 (TLR4) (yellow) (Fig. 4d-e, Fig. 5b-g).

Confocal microscopy of the Sham group revealed low basal expression of both markers, in the CPE, the intact ependymal lining and the cortex (Fig. 4d, e, Fig. 5b, d, k). TLR4 was also distinctly expressed in the blood vessels of all groups (white solid arrows) (Fig. 4d and Supplementary Fig. 2d). In contrast, untreated PHH-Control demonstrated robust NLRP3 and TLR4 expression (dotted white arrow) (Fig. 4d, e), along the ependymal border and extending into the underlying anterior subventricular zone (ASVZ) parenchyma, corresponding to regions of structural disruption. High expression of TLR4 in the ependymal cilia was also observed (white arrow) (Fig. 5h). Treatment with GSK2789917 in the PHH-Treatment group visibly attenuated NLRP3 fluorescence in some areas of the CPE (white arrow) (Fig. 4d, e).

Quantification of fluorescence intensity within the ependymal lining (Fig. 5i, j), revealed a significant elevation in NLRP3 expression in the untreated PHH-Control group compared to Sham controls (p < 0.05). There was a highly significant reduction in NLRP3 intensity in PHH-Treatment compared to PHH-Control (p < 0.001), restoring levels to values not significantly different from Sham (Fig. 5i). Similarly, TLR4 fluorescence intensity was significantly elevated in PHH-Control compared to Sham controls (p < 0.05). Treatment significantly reduced NLRP3 intensity in the CP compared to PHH-Control (p < 0.001) and even the Sham-operated group (p < 0.05) (Fig. 4b). There were no significant differences in TLR4 intensity between the Sham, PHH-Control, or PHH-Treatment groups in the CP (Fig. 4c), although some areas of the PHH-Control CPE showed high expression (Supplementary Fig. 2e).

To further explore these neuroinflammatory markers in the CPE, we performed linear regression analysis. This revealed a significant positive correlation between TLR4 and NLRP3 fluorescence intensity across individual samples (R² = 0.366, F (1,31) = 17.89, p = 0.001) (Fig. 4g). These findings indicate that increased TLR4 expression is associated with elevated NLRP3 protein expression within the tissue, particularly in samples from the untreated PHH-Control group (dotted white arrows) (Fig. 4d–f; Supplementary Fig. 2e).

Treatment with GSK2789917 attenuated NLRP3 fluorescence intensity within the periventricular region (Fig. 5b). Within the periventricular WM (Fig. 5e; Supplementary Fig. 2a), NLRP3 expression was significantly elevated in the untreated PHH-Control group compared to Sham controls (p < 0.05). The cerebral cortex (CTX) exhibited robust NLRP3 expressions characterized by increased density and intensity of positive cells throughout the cortical field (Fig. 5f, g). There was a significant elevation in cortical NLRP3 fluorescence intensity in untreated PHH compared to Sham controls (p < 0.01) (Fig. 5f), accompanied by a significant increase in the proportion of cortex cells expressing NLRP3 (p < 0.001) (Fig. 5g). Treatment with GSK2789917 significantly reduced both NLRP3 intensity (p < 0.01) and proportion (p < 0.01) relative to untreated PHH. Notably, both metrics were restored to levels not significantly different from Sham controls (Fig. 5f, g). TLR4 fluorescence intensity within the CTX was visually increased in untreated PHH compared to Sham and treated groups (white arrow) (Fig. 5k).

## DISCUSSION

Hydrocephalus remains a complex, debilitating and life-limiting disease, compounded by the lack of non-surgical therapy at the clinical level and by insufficient modeling at the pre-clinical level. In this study we established that GSK2789917 traverses the BBB and NVU. We demonstrated that the compound possessed the oral bioavailability predicted to reach efficacious concentration in pigs. Exposure did not appear to be affected by induction of PHH. The TRPV4 antagonist decreased pathological parameters of hydrocephalus including ventriculomegaly, increased ICP, and neuroinflammation. These results establish a valid, gyrencephalic, severe *in vivo* model and provide compelling justification for human TRPV4 antagonist clinical trials.

### A severe and sustained swine PHH model

The newly developed swine model incorporates intraventricular autologous blood mixed with inert embolism beads, mimicking the cellular debris of IVH. Importantly, the model does not require caustic chemicals such as kaolin and does not rely on exogenous cellular toxicity.

The model is severe, with a challenging anesthesia recovery and an early fatality rate of 23.5% (4/17 pigs with accurate LV injection succumbed during the first night). The disease induction was comparable to a severe trauma or intraventricular bleed, with fatally high intra-operative ICPs, and neurological impairment of all surviving pigs. While life-threatening neurological impairment did not extend > 24 hours, it should be noted that pigs are not dependent on a functioning cerebral motor cortex to ambulate. Instead, pigs with a severe hemispheric lesion can ambulate, circling compulsively towards the side of the more severe lesion, as observed in all pigs.

The validity of the model is best informed by examining the difference between the Sham-operated and PHH-Control groups. The Total Ventricular Volume was significantly larger in the PHH-Control group on all post-operative scans (Days 10 – 24), indicating a sustained and marked ventriculomegaly, predominately due to increases in the lateral ventricles (Fig. 2).

Enlargement of the 3^rd^ and 4^th^ ventricles did not develop. This indicates an obstructive, non-communicating hydrocephalus, with enlargement limited to the lateral ventricles. While a fraction of the blood-bead mixture did enter the 3^rd^ ventricle, it did not obstruct passage of CSF into the aqueduct. The intraventricular hematoma resolved (extended data Fig. 1c), but ventriculomegaly persisted throughout the study (Fig. 2b), as IVH evolved into PHH.

Congruent evidence of model validity was evident in the ICP results. For the Sham operated pigs, mean ICP remained in the normal range (<20 mmHg) throughout the study. There was an initial, small upwards trend, coinciding with opioid analgesia. From Day 5, ICP trended downwards, returning to baseline. In contrast, the mean PHH-Control ICP trended steeply upwards until Day 10; from Day 3 – 11, the mean ICP was in the elevated range (20 – 40 mmHg). The ICP-time relationships of all 3 groups were different (P < 0.0001), and *post hoc* the PHH-Control ICP was significantly (P < 0.05) higher than the Sham group by Day 10. The ICP and Total Ventricular Volume data matched, in both the PHH-Control pigs (peaking at Day 10, then decreasing for the remainder of the study) and the Sham group (no major changes in either parameter).

Post-mortem histology and immunofluorescence revealed further powerful support of severe and persistent PHH. While the hematoma had resolved, ventriculomegaly was still present histologically, upon sacrifice 30 days after surgery. Sham animals exhibited preservation of the CP architecture, ependymal lining and subventricular white matter (Figs. 4, 5 and Supplemental 2). In stark contrast, the PHH-Control group displayed ependymal disruption, stromal edema and vascular dilation within the CP, periventricular WM degeneration, and cortical neuronal vacuolation.

Further, Sham animals displayed only basal expression of the inflammatory markers NLRP3 and TLR4. Compared to Sham, NLRP3 fluorescence was significantly elevated in the ependyma, white matter and cortex of PHH-Control pigs. TLR4 showed similar patterns, with significantly increased expression in the ependyma. The intensity of NLRP3 and TLR4 in the CP were significantly correlated. The neuroinflammatory marker findings support that the PHH induction caused inflammation from the ependyma to the subcortical white matter.

In summary, this swine model demonstrates a severe and sustained post-hemorrhagic hydrocephalus with many correlates to human disease.

### Control of ventricular size and ICP

GSK2789917, begun 2 days after the hemorrhagic insult, strongly inhibited ventriculomegaly (PHH-Treatment *vs* Control, Fig. 2). Strikingly, this effect was maintained for a week after discontinuing drug treatment. This suggests that if TRPV4 antagonism treatment begins in a timely fashion after IVH, hydrocephalus might be prevented. This potentially relates to a decrease in CSF production by choroid plexus cells.^5^ Supporting reduced CSF production, the PHH-Treatment group had significantly smaller 3^rd^ and 4^th^ ventricles (Fig. 2c) than the Sham on all post-treatment MRIs, and smaller 4^th^ ventricles than the Control pigs (Days 17 and 24).

The ICP results from the PHH-Control and PHH-Treated group also support a considerable beneficial impact of TRPV4 antagonism (Fig. 3). The baseline Day 1 – 2 values were similar, around 10 mmHg as previously observed.^7^ GSK2789917 therapy began on Day 3, coinciding with a considerable divergence between the two PHH groups. The PHH-Control ICP continued to trend upwards for the next week, spending a substantial amount of time in the elevated (20 – 40 mmHg) zone. The interaction between treatment group and time was significant (P < 0.0001), indicating that ICP trends differed between groups. Additionally, in *post-hoc* tests the difference between Control and Treatment groups reached significance on Day 10 (Fig. 3c). In the Treatment group, ICP remained largely static during the first week. There was then an upwards trend during the remaining period of GSK2789917 administration. A possible explanation is that unmitigated CSF production forces recanalization through intraventricular hematomas^7^, a process which is delayed by pharmacological reduction of CSF production. This would allow a slow increase in ICP, as CSF accumulates over multiple days. The highest mean ICP in the Treatment group was in fact seen the day after discontinuation, after which ICP values fell towards baseline. In contrast, the Control group showed an initial rapid increase in ICP, followed by a drop in ICP that might coincide with recanalization. Additionally, there is a possible impact of increasing brain health leading to increased movement artifact and higher apparent ICPs.

Strikingly, comparison of the PHH-Treatment and Sham groups suggest that GSK2789917 might reverse components of IVH. Certain metrics were the same or even lower, in the Treatment group. This included no significant difference in post-operative Total Ventricular Volumes (Fig. 3d), significantly *smaller* 3^rd^ and 4^th^ ventricles (Fig. 3c), significantly *lower* ICP on the first 2 days after beginning GSK2789917 (Fig. 3d), and finally lower NLRP3 in the CP (Fig. 5b).

### GSK2789917 reduces neuroinflammation

The present model of PHH reveals widespread histopathological alterations in the choroid plexus, ventricular lining and brain parenchyma, accompanied by increased expression of damage-associated molecular patterns marker TLR4 and activation of NLRP3-dependent inflammasome signaling. These findings indicate a close association between structural disruption and inflammation activation. Notably, pharmacological treatment with GSK2789917 attenuates both tissue injury and activation of the TLR4–NLRP3 inflammatory axis. Tissues were collected for analysis at Day 31, two weeks after the last dose of the TRPV4 antagonist, indicating a sustained beneficial effect.

Within the CP, PHH-induced architectural alterations are accompanied elevated NLRP3 expression in epithelial cells. These findings are consistent with rat IVH models, in which NLRP3 inflammasome activation peaks early and localizes to the CPE.^16^ Mechanistically, NLRP3 activation in this compartment is linked to mitochondrial dysfunction, increased reactive oxygen species production, and disruption of epithelial tight junctions, compromising blood–CSF barrier integrity.^17^ In the present study, treatment with a TRPV4 antagonist attenuated both the structural abnormalities and NLRP3 expression, suggesting that reduced inflammasomes is not solely a direct pharmacological effect but may also reflect the mitigation of the underlying hemorrhage-induced microenvironmental stressors that drive inflammasome activation.

PHH also resulted in severe disruption of the ependymal lining and inflammatory activation that extended into the ASVZ. TLR4 and NLRP3 expression were significantly elevated along the damaged ependymal surface, consistent with structural disruption acting as a potent trigger for inflammation. GSK2789917 ameliorated the ependymal damage and NLRP3 expression, supporting a mechanistic link between structural preservation and suppression of inflammasome activation. These observations align with ependymal damage, a 75% reduction in SVZ proliferative cells, and neuroinflammation within the periventricular WM and CSF, in kaolin-induced hydrocephalic pigs.^18^ Human fetal studies of hydrocephalus similarly report ependymal denudation progressing with gestational age.^19^

This structure–inflammation coupling extends into the periventricular WM and CTX, which are particularly vulnerable in PHH-induced ventriculomegaly. Animals in the PHH-Control cohort exhibited marked WM disorganization with increased NLRP3 and TLR4 expression, consistent with studies linking hydrocephalus to white matter injury and inflammation.^18,20^ In the cortex, PHH induced cell vacuolation and significant NLRP3 expression. Treatment with GSK2789917 effectively reduced these structural integrity and inflammasome alterations in the WM and CTX. These findings reveal that the hemorrhage-induced cellular injury initiates a propagating inflammatory cascade that extends beyond the ventricular interface into distant parenchymal regions, and that interruption of this cascade with TRPV4 antagonist treatment limits widespread neuroinflammatory engagement.

Collectively, these MRI, histology and immunofluorescence data support a hydrocephalus model in which hemorrhage-induced structural distortion triggers the activation of inflammation across multiple neuroanatomical compartments. Treatment with a TRPV4 antagonist mitigates the structural alterations and the ensuing neuroinflammation, offering a comprehensive approach to preserving brain integrity.

### Translating TRPV4 inhibition into humans

TRPV4 antagonists have been evaluated in humans for a variety of indications. GSK2789917, provided by GSK, is a TRPV4 inhibitor with >100-fold selectivity for human TRPV4 (IC_50_ 3nM) compared to more than 100 other molecular targets.^21^ Being from the same benzimidazole spirocarbamate chemical series, it shares multiple structural and functional characteristics with TRPV4 inhibitor GSK2798745, which was recently assessed in multiple cardiorespiratory clinical trials with no adverse side effects.^8–12^ These studies included healthy volunteers^11^, stable heart failure patients^8^, chronic, compensated heart failure patients^10^ and refractory chronic cough patients.^12^ There was no mortality or serious drug-related adverse effects, and GSK2798745 was well-tolerated by cardiorespiratory patients and healthy participants.^8–12^ Actio Biosciences is conducting a Phase 1 trial for a small molecule TRPV4 inhibitor, ABS-0871, with the targeted indications of Charcot-Marie-tooth 2C Syndrome and overactive bladder.

In summary, clinical trials have indicated that TRPV4 antagonists are safe, well-tolerated and display no reported adverse effects. This is consistent with the finding that *Trpv4* null mice have a normal appearance, growth and reproductive capacity, and only develop osmotic abnormalities if placed under severe osmotic stress.^22^ Combined, these data suggest that inhibition of TRPV4 as a clinical therapy will be safe.

In the current studies, the benefit of the TRPV4 antagonist on the ventriculomegaly persisted on the MRI 7 days after treatment was terminated, while the reduced structural and inflammatory changes were seen 14 days after the last dose, indicating long-lasting beneficial effects. These compelling results indicate that TRPV4 antagonists are effective in a large animal, gyrencephalic model of PHH. Combined with the rodent data suggesting that multiple TRPV4 antagonists are effective in a genetic model of hydrocephalus^6^ and in PHH^23^, our results provide an enhanced indication for the success of TRPV4 inhibition in treating multiple forms of hydrocephalus.

### Limitations

The main limitations in this study are reduced signal-to-noise ratio for the ICP recordings, necessitating exclusion of ICP data from some subjects, combined with limited final cohort sizes and, therefore, male-only pigs.

During surgery, telemetric epidural ICP recordings matched the gold standard (intraventricular ICP, performed intra-operatively in this study). Both epidural and intraventricular ICP traces had small, rapid waves matching the cardiac cycle, with larger, slower waves matching respiration (Fig. 3a), and immediate increases upon blood injection. Interpretation of the ICP recordings on awake pigs was more complex. The pressure sensor / battery pack unit was implanted over the neck, to prevent MRI artifact superimposed over the brain. This unit was designed to be placed adjacent to the rodent brain.

In a large mammal such as a pig, raising and lowering the head will change the relative position of the probe (inside the skull) to the sensor by multiple centimeters. This artifactually changes the ICP value. Head motion also changes the true ICP, increasing as the head is lowered.

In sleeping post-operative animals, traces remained easily interpretable, including very small waves matching the cardiac cycle. However, as soon as pigs began to move, movement artifact created far larger deflections. During data analysis, these movement artifact waveforms were easily recognized and excluded, due to their extremely rapid and enormously large deflections. It is important to note the possible bias this introduces. Excluding pigs with excessive movement artifact likely removed the more active pigs.

The limited, male-only cohort sizes were due to financial constraints, exacerbated by considerable study dropout (8/21, 38%). The severity of the model, in which multiple accurately injected subjects died within 24 hours, accounted for half of the dropout. Further, a PHH-Control pig succumbed to recrudescence of neurological symptoms before the final MRI. With such small cohort sizes, the loss of the subject with the largest ventricles might have diminished the Day 24 mean Total Ventricular Volume (Fig. 2a).

## METHODS

### *In vitro* BBB and NVU permeability studies

To mimic the *in vivo* BBB, we have developed and optimized a novel, layered direct contact triculture (human astrocytes, pericytes and brain microvessel endothelial cells) Transwell system.^14^ Enhancement of this model was made by co-culturing the neuronal cell line, SH-SY5Y, with the BBB triculture cells to mimic the *in vivo* NVU.^13^ Bidirectional permeability studies were performed as described^13–15^, to assess the efflux ratio [ER, (P_APP Bl→Ap_)/(P_APP Ap→Bl_)] to determine the relative exposure rates at steady state (SS). Furthermore, at C_SS_, the K_p,uu_ is defined as the rates into the CNS divided by the outward rate of the CNS, or CNS_in_/CNS_out_. Thus, the 1/ER approximates the therapeutic effects in the brain parenchyma estimated by the K_p,uu_ at steady state. Levels of GSK2789917 were quantitated using a gradient high-performance liquid chromatography method on an Agilent 1100 HPLC.

### GSK2789917

GSK provided GSK2789917, (C, (1-(((5S,7S)-3-(4,6-dimethoxypyridin-3-yl)-7-methyl-2-oxo-1-oxa-3-azaspiro[4.5]decan-7-yl)methyl)-1H-benzo[d]imidazole-6-carbonitrile), under a Material Transfer Agreement; animal work described in this publication was not conducted on behalf of GSK.

### *In vivo* pharmacokinetic studies

#### Single Dose Pharmacokinetic Parameters – non-PHH pigs

Pigs were socialized for 7 days. A surgical procedure was performed 24 hours before the pharmacokinetic study to insert the jugular catheter, externalized in the dorsal scapular region. Pigs were then placed in a movement responsive cage system (Culex-L^®^, Bioanalytical Systems Inc., West Lafayette, IN, USA) to prevent entanglement of the catheter lines. When the animal rotates more than 270°, the system receives a signal transferred through the catheter to respond with a cage movement in the opposite direction. This facilitates automated blood sampling and minimizes stress.^1–3^ Briefly, the drug was portioned into size 0 capsules, at 60 mg per capsule, to provide a dose of approximately 2 mg/kg in the *circa* 30 kg pigs. Two capsules were administered to create 4 mg/kg doses. Blood samples of 1 mL were collected at 0.25, 0.5, 1, 1.5, 4, 6, 8 and 12 hours after dosing and transferred into vials containing tri-potassium ethylenediaminetetraacetic acid. The vials were centrifuged at 4 °C and processed to extract plasma, which was then stored at –80 °C until liquid chromatography-tandem mass spectroscopy analysis. An Agilent 1200 Rapid Resolution LC system coupled to an Agilent 6460 series QQQ mass spectrometer was utilized. An Agilent PFP 2.1 mm x 100 mm, 3 µm column was used for liquid chromatography separation (Agilent Technologies, Santa Clara, CA). The method was adapted, validated, and then used to analyze samples extracted from porcine plasma. Multiple reaction monitoring was used for MS analysis. All data were analyzed using Agilent MassHunter Quantitative analysis software (version 6.0). The calibration curve range was 10,000-1 ng/mL in plasma. The correlation coefficient (r^2^) was >0.999 and all data were normalized to the ^13^C_2_, ^15^N_1_-internal standard peak.

#### Multiple Dose Pharmacokinetic Parameters – non-PHH pigs and PHH pigs

We performed multiple dosing studies with GSK2789917 at a dose of 2 mg/kg two times a day, to evaluate exposure in normal pigs, and to determine accumulation and the relationship to efficacy in the hydrocephalic model. Blood samples were collected (n = 4 normal males) at the times in the single dose study above, with an additional sample 24 hours after dosing (days 1, 5, and 8). Blood was also collected 2, 12, and 24 hours after dosing on the intermediate days. After collection, samples were analyzed as above.

A steady-state plasma concentration (C_p_,ss) was not estimated, as this would have required additional plasma samples, adding additional stress to the animals. The determined maximum concentration (C_MAX_) on days 5 and 8 reveal that levels were above our predicted Effective Concentration 100% (EC_100_; the concentration required to produce 100% of the maximum biological effect in the brain) throughout the studies. The plasma profiles appeared to be approaching C_p,SS_ levels, although the study period was not sufficient to determine if linear pharmacokinetics would be achieved. The T_max_ values were not significantly different across the days, except for one outlier at day 5. The AUC values reflect accumulation, with an apparent increase in the half-lives (T_1/2_) over time. The apparent increase in T_1/2_ was reflected in a decrease in the elimination rate, although the results between days 5 and 8 were not significant.

During the subsequent *in vivo* surgical study, blood samples were collected from the PHH-Treatment group (n = 6), starting on Day 3, following the first dose of GSK2789917. These samples were analyzed as a separate multiple dose PK study. Due to two of the pigs experiencing seizures after the surgery, palliative therapies including sedation were provided, which we postulate slowed absorption. The plasma concentration *vs.* time curves from these two pigs differed significantly from the other four and were not included in the final PK parameter analysis reported here. Blood samples were collected as described above, 0.25 to 24 hours after dosing, including an additional sample 2 hours after dosing.

#### Pharmacokinetic Analysis

Data fitting and parameter estimation of GSK2789917 concentration-time data were performed using the non-compartmental model extravascular analysis on for the first dose on Days 1, 5 and 8 in the PKMP software from APL Analytics (Eden Prairie, MN, USA). The PK parameters investigated include maximum plasma concentration (C_max_), time to reach the maximum concentration (T_max_), area under the concentration-time curve (AUC), half-life (T_1/2_), elimination rate (Kel), Mean Residence Time (MRT), volume of distribution (V_d_), and clearance (Cl). A one-compartment model with first-order elimination was used to simulate the data.

### Pig cohorts

Barrows from Oak Hill Genetics (Ewing, IL) were housed in accordance with protocol 0223002360. Animals were maintained on a 12-hour light cycle with Envigo 7037 swine diet (Indianapolis, IN) daily and ad lib water. Animals were fasted prior to the 24-hour PK study and fed 2 hours after dosing with GSK2789917.

Four cohorts of pigs were utilized. The first cohort was used for pharmacokinetic studies only (4 pigs per study). The other 3 cohorts underwent Sham surgery (n = 4) or IVH surgery (n = 21). Pigs who survived at least 24 hours after IVH surgery and were confirmed to have accurate intraventricular injection of blood (n = 13) were randomized into PHH-Treatment (n = 6) and PHH-Control (n = 7) cohorts.

Following surgery, pigs were removed from the study and euthanized if they did not regain ambulation within 3 hours. All pigs displaying severe neurological disability impairment (persistent recumbency and vestibular syndrome despite 1 gr/kg mannitol IV; recurrent seizures despite 0.5 mg/kg diazepam IV per seizure) were sacrificed and removed. A total of 21 pigs underwent a right LV injection of 12 ml of a solution containing 10 ml of autologous blood and 2 ml of 700 µm embolism beads. 8 of these pigs were removed from the study (see below).

### Surgical study

#### MRI

MRI was performed immediately prior to the surgery (Day 1), and repeated on Day 10 ± 1, Day 17 ± 1 and Day 24 ± 2. In the PHH-Treatment group, GSK2789917 was administered on Days 3 – 17. Post-operative MRI thus occurred after 7 and 14 days of GSK2789917, and 7 days after discontinuing it. A 3 Tesla Siemens MAGNETOM Prisma MRI scanner was used. T2-weighted (T2W), T2-SPACE ISO and T1-MPRAGE sequences were acquired in dorsal, sagittal and transverse planes. T1-weighted (T1W), T2*-weighted and T2W FLAIR images were acquired in transverse planes. Semi-automated measurement of ventricular size was performed with Imalytics software, utilizing the T2-weighted MRI sequence. The outline of the ventricles was then adjusted as necessary for accuracy on each individual slice by FOB. The 3-dimensional volume of each ventricle was then calculated (1 voxel = 0.80719 mm^3^). Each of the following ventricles was measured separately: LV (left and right), olfactory recess of the LV (left and right), 3^rd^ ventricle and 4^th^ ventricle. The combined volume of these ventricles was the Total Ventricular Volume. Ventricular size analysis was performed by two investigators blinded to the study design.

#### CT scanning

For the final 12 IVH pigs, a CT was performed within 24 hr of surgery, to rapidly confirm accurate intraventricular injection of blood. CT scans were performed using an 80-slice multi-detector CT scanner (Aquilion PRIME; Canon, Tokyo, Japan). Parameters included a helical scan mode, pitch of 1.388, 0.5 s rotation time, 120 kVp, 350 mA, and 0.5 mm slice thickness. No results from pigs with extra-ventricular blood injection on CT or MRI were included in the study.

#### Surgical methodology (Day 1)

Pigs were premedicated with midazolam (0.2 mg/kg IM) and dexmedetomidine (0.03 mg/kg IM), then anesthetized with 5% isoflurane in oxygen via facemask. Endotracheal intubation (5.5 - 6 mm) was performed and pigs were maintained on 1-2% isoflurane in medical oxygen. Antibiosis included ceftiofur (5 mg/kg IM, pre-operative depot formulation, minimum 4-day activity) and florfenicol (16.5 mg/kg IM, given twice, before and after surgery). Analgesia included intra-operative fentanyl (10 mcg/kg/hr), and post-operative buprenorphine (0.03 mg/kg BID, IV or SQ x 72 hr). Intra-operative monitoring included ECG, capnography, temperature and direct arterial pressure measurement. Pigs received 5 ml/kg/hr lactated Ringer’s solution and active heat support. During surgery, a mechanical ventilator was used to maintain EtCO2 at a target of 35 mmHg.

Pigs were anesthetized, the region of the left jugular was prepped and draped for sterile surgery, and a left jugular catheter was placed, and externalized in the left dorsal scapular region. The dorsal head, neck and interscapular region was prepped and draped for sterile surgery. A curving skin incision was made over the skull (external occipital protuberance to the right zygomatic process). The right temporalis muscle fascia was incised and the right temporalis muscle was elevated. A burrhole craniectomy was made, roughly 1 cm height x 2 cm length, in the right frontal bone, immediately caudal to the zygomatic process. An 18G intravenous cannula was placed through the intact dura mater into the right lateral ventricle. The correct ventricular location of the cannula was confirmed by the passage of CSF. This cannula was connected to sterile non-compressible tubing, permitting intra-operative measurement of the intraventricular ICP. A second cannula was placed immediately adjacent, and the ventricular location again confirmed by the passage of CSF. This cannula was used to perform the intraventricular blood injection.

Another incision was made through the skin and subcutaneous muscle, on the dorsal neck, cranial to the right scapula. A pocket was created under the subcutaneous muscle. A trocar was passed through the right temporalis muscle into the pocket of dead space. The probe of an implantable telemetric epidural ICP monitor (Data Sciences International) was passed through the trocar. The recording tip of the probe was introduced into epidural space (between the dura mater and the frontal bone). The battery pack was placed within the subcutaneous pocket. The trace of both ICP measurements (intraventricular and epidural) were visually confirmed to match the cardiac cycle (upswing with systole and downswing with diastole), and to increase with each inflation of the thorax by the mechanical ventilator. The intraventricular ICP was confirmed to have a baseline of 9 – 12 mmHg; the reading from the epidural ICP probe was more variable and would typically be slightly higher, due to the battery back being gravitationally above the recording tip of the probe.

Blood (total volume, 10.0 mL) was drawn from the jugular catheter in 3 separate aliquots of 3.33 mL. The syringes were handled by an assistant wearing sterile surgical gloves and passed to the sterile surgeon. A total volume of 2.0 mL of endovascular embolic beads (Embozene; 700 µm diameter) (Varian Medical Systems INC, Austin, TX) was added, by adding one-third of the beads to each syringe. The total volume injected was therefore 12.0 mL. During injection of the first 4.0 mL syringe, measurements from both ICP probes would immediately increase, then rapidly trend down again. The intraventricular ICP remained under 50 mmHg for the entire injection. The second syringe was then injected, pausing each time the intraventricular ICP exceeded 50 mmHg, after which it would trend downwards. The final syringe was not drawn until the ICP was under 50 mmHg. During injection of the 3^rd^ syringe, elevation of the intraventricular ICP above 50 mmHg was common, and it would trend back down quite slowly. It therefore took 2 – 5 minutes to inject this syringe. After the injection was completed, it would take another 2 – 5 minutes for the intraventricular ICP to fall below 50 mmHg. Both intraventricular cannulas were then removed.

The temporalis fascia was closed with 3-0 PDS, immobilizing the probe of the ICP monitor in the epidural space. The subcutaneous tissue and skin of the cranial surgical site was closed in 2 layers of poliglecaprone, followed by skin glue. The battery pack surgical site was closed in 2 layers of poliglecaprone, followed by skin glue.

### Post-operative monitoring and care

The jugular catheter was left in place for post-operative medications and PK analysis. Most pigs regained ambulatory status and began eating and drinking within 3 hours. Seizures were treated with 0.5 mg/kg diazepam IV. Animals with severe neurological deficits (e.g. remaining non-ambulatory more than 3 hours after the end of anesthesia) were euthanized and removed from the study.

Pigs were observed continuously by a board-certified veterinary neurologist during anesthesia recovery, and re-examined twice a day on Days 2 – 4. Examination included mentation, gait, posture, cranial nerve deficits and proprioceptive responses. A semi-quantitative score for neurological deficits was developed (Table 1).

Surgery was identical for PHH-Treatment and PHH-Control group pigs. Surgeons were blinded to the group: pigs were allocated to Treatment *versus* Control post-operatively. For the Sham surgery group (n = 4), the surgical approach and closure was the same as above. The implantable epidural ICP monitor was implanted as above. No cannulas were placed into the lateral ventricles and no material was injected.

The study was approved by the Purdue Institutional Animal Care and Use Committee (Protocol 0223002360).

### Adverse events and exclusions

#### Pigs removed from the study

Of the 21 injected pigs, 8 were removed from the study. Two were euthanized because of a severe post-operative neurological condition (remaining non-ambulatory and obtunded > 3 hours post-anesthesia, combined with recurrent seizures in one pig). Another 2 pigs regained ambulation post-operatively but were found dead the next morning. Both had displayed mania during their initial anesthesia recovery, compulsively running into walls, attenuating over the first hours.

There were 4 pigs removed from the study because blood was not accurately injected into the LV (blood located superficially in the parenchyma of the right hemisphere (n = 3), or in the quadrigeminal cistern (n = 1)). For the first 9 surgeries, the Day-10 MRI was evaluated for correct intraventricular injection of blood. For the remaining 12 pigs, a CT was performed within 24 hr of surgery, allowing exclusion of incorrectly injected pigs prior to allocation to PHH-Control *vs* PHH-Treatment groups.

#### Pigs maintained in the study – adverse events

6 of the 13 pigs had between 2 and 4 seizures, during the first 4 hours after surgery. One additionally showed temporary vestibular dysfunction. Each seizure was treated with 0.5 mg/kg diazepam IV. One pig became sedated due to repeated diazepam and received IV fluid therapy overnight to maintain hydration. All 6 pigs were ambulatory and eating the morning after surgery, had no more seizures and were maintained in the study.

On Day 3 (when medication was started in the PHH-Treatment group), all pigs were well, seizure-free (> 36 hours), and were neurologically stable and improving.

### Allocation to Control *vs* Treatment

Pigs alive 24 hour post-operatively were allocated to the PHH-Control group (n = 7) or the PHH-Treatment group (n = 6). (While 7 pigs were initially started on GSK2789917, in one pig failure to accurately inject the LV was identified on the Day 10 MRI. All data from this animal were excluded.)

#### Administration of TRPV4 Antagonist

On Days 3 – 17, GSK2789917 was administered to the Treatment group at a dose of 60 mg (*circa* 2 mg/kg), PO BID.

### Neurological progression of pigs

In the PHH-induced pigs, upon beginning to wean the mechanical ventilator during surgical closure, it took 15 to 30 minutes for spontaneous respiration, and 1 – 2 hours for extubation.

For 2 – 6 hours after extubation, pigs showed small improvements as anesthesia diminished. All 13 pigs initially had variable reduced +/-delirious mentation, reduced-to-absent vision, a head turn and circling to the right, paresis and ataxia. As anesthesia resolved, pigs returned to ambulation, being particularly ataxic and paretic in the pelvic limbs. They had to be restrained to prevent compulsively circling to the right and falling. All pigs were blind in their left visual field; while circling, they would bump the left face into cage bars. Vision improved variably during the hours following anesthesia. Some pigs regained vision initially, but were temporarily blind again post-ictally.

By 6 hours post-extubation, the menace response was absent in the left eye (100% of pigs) and reduced-absent in the right eye (9/13, 69%). Most pigs were ambulatory, eating and drinking, with proprioceptive deficits in all 4 limbs (worse on left). Individual pigs showed additional deficits: seizures (n = 6), head-pressing after diazepam to treat seizures (n = 4), persistent right head turn and circling (n = 5), persistent paresis ± ataxia with falling (n = 2), slowly resolving recumbency (n = 2), mild mania (without self-trauma, allowing sedation rather than euthanasia, n = 1), tremors (n = 1) and resolving vestibular syndrome (regaining ambulation within 3 hours, n = 1).

During the first 3 days after surgery, pigs were awarded a semi-quantitative neurological score based on neurological examination by a board-certified veterinary neurologist (Table 1). On Day 2, severity varied from one pig with only a unilaterally reduced menace response, to another with 7 neurological deficits (reduced mentation, circling to the right, ataxia, bilaterally menace response deficits, bilateral proprioceptive deficits). By Day 4, every pig showed some combination of reduced-absent menace response OS (all pigs), turning right more often than left (n = 10), reduced menace response OD (n = 3), and mild ataxia without falling (n = 2), with no other neurological deficits.

One pig displayed 6 neurological deficits including pronounced lethargy on Day 2 (Table 1). Showing typical improvement after surgery, by Day 4 he was bright with only 2 neurological deficits. On Day 7, unlike all other pigs, he was dull again for one day. On Day 22, he was lethargic again, inappetent, began having seizures and became hyperthermic. He was sacrificed, without performing the Day 24 MRI. His data were maintained in the study. This pig had been allocated to the PHH-Control group, on Day 2 as per study protocol.

Every Sham-operated pig was neurologically normal by 1.5 hr post-extubation, and none suffered any seizures. During Days 2 – 4, every Sham pig remained neurologically normal, apart from one mildly reduced menace response on Day 2 (Table 1).

### Epidural ICP processing

Any material (such as surgical debris or hemorrhage) entering the tip of the sensor dampens recordings. Dampening can be detected by noting decreased deflections associated with the cardiac and respiratory cycles. If this was noted intra-operatively, the tip of the sensor was removed from the epidural space, cleaned, refilled with gel, and re-implanted.

The epidural ICP measurements were processed in Ponemah software, analyzed using Python, and visualized in GraphPad Prism 8.0. Movement artifact was excluded from the automated daily mean ICP calculation with a 30 Hz Low Pass Filter, and by excluding values that were artifactually high (>80 mmHg) or low (<0 mm Hg) or had a frequency incompatible with the cardiac cycle (< 30/min or > 300/min). Only subjects with adequate ICP data were included in the automated calculation of the daily mean ICP for the 3 cohorts (PHH-Control, PHH-Treatment, Sham).

### Histology and immunofluorescence

Following euthanasia, the head was removed at the atlanto-occipital joint. A handsaw was used to perform circumferential craniotomy. The dorsal surface of the brain was exposed, and the dura mater was carefully incised. The brain was gently freed from the cranial floor by sequential transection of cranial nerves and the optic chiasm. The intact brain was removed, hemi-sected, and one half was placed in 10% neutral buffered formalin for post-fixation and then processed and embedded in paraffin according to standard procedures.

Brain removal allows the release of CSF and the collapse of enlarged ventricles. *Ante-mortem* MRI, showing significantly increased ventricular size in PHH-Control pigs (Fig. 2) provided a more complete representation of ventriculomegaly. Despite this, ventricular size was enlarged in PHH-Control pigs upon histological examination (Fig. 5a), corroborated by *ante-mortem* ependymal stretching and disruption.

#### Hematoxylin and Eosin (H&E) Staining

General tissue morphology was assessed using an H&E staining kit (Novus Biologicals, Cat# H-3502-NB). Paraffin-embedded sections (4 µm) were deparaffinized in xylene (two changes, 5 min each) and rehydrated through a graded ethanol series (100%, 95%, and 70%), followed by rinsing in DDI water. Sections were incubated in Mayer’s hematoxylin for 5 min to visualize nuclei, rinsed twice in distilled water, and differentiated using a bluing reagent for 15 s. After two additional washes in distilled water, slides were immersed in 100% ethanol (10 s) and counterstained with Eosin Y solution for 3 min to label cytoplasmic and extracellular components. Following staining, sections were rinsed in 100% ethanol, dehydrated through three changes of 100% ethanol, cleared in xylene, and coverslipped using a permanent mounting medium.

#### Immunofluorescence Staining

Paraffin embedded tissue sections (5 µm) were deparaffinized in xylene (two changes, 5 min each) and rehydrated through a graded ethanol series (100%, 95%, and 70%), followed by rinsing in DDI water. Followed by antigen retrieval, which was performed by heating sections in 10 mM sodium citrate buffer (pH 6.0; 0.05% Tween-20) to boil for 10 min, slides were allowed to cool for 30 min at room temperature while submerged in retrieval buffer, then washed in PBS. To reduce tissue autofluorescence, sections were treated with TrueBlack® (Biotium) Cat 23007. A 1× working solution was freshly prepared in 70% ethanol, and 100 µL was applied to each section for 2 min at room temperature. Sections were then rinsed three times in PBS. Sections were blocked for 1 h at room temperature in blocking buffer (1% BSA in PBS). Sections were incubated overnight at 4°C with primary antibodies diluted in blocking buffer. For co-localization studies, primary antibodies raised in different host species were applied simultaneously. Primary antibodies included anti-TLR4 (mouse, 1:100; Abcam, ab22048) and anti-NLRP3 (rabbit, 1:100; Novus Biologicals, NBP2-12446SS). After washing in PBS, sections were incubated for 1 h at room temperature with species-specific secondary antibodies: Alexa Fluor 488 goat anti-mouse IgG (1:1,000; Abcam, ab150113) and Alexa Fluor 546 goat anti-rabbit IgG (1:1,000; Invitrogen, A-11035). Nuclei were counterstained with 4′,6-diamidino-2-phenylindole (DAPI; Sigma-Aldrich, D9542). Following final PBS washes, sections were mounted with an aqueous antifade mounting medium (Aqua-Poly/Mount, Polysciences, Inc.), coverslipped, and stored protected from light until imaging.

### Statistical Analyses

#### MRI

Statistical analyses were performed in GraphPad Prism using a two-way ANOVA to assess group and time effects and applied a mixed-effects model where there were repeated measurements from the same animals. Asterisks indicate statistical significance based on Tukey-corrected multiple comparisons within each time point.

#### ICP

Statistical analyses included a mixed effect model post-hoc multiple t-tests (Holm’s correction).

#### Immunofluorescence

Statistical analyses were performed using GraphPad Prism (version 10). Data are presented as mean ± Standard Error of the Mean (SEM): n = 3 (Sham), n = 4 (PHH-Control), and n = 4 (PHH-Treatment). 3 distinct aspects of the same tissue per animal were analyzed for CPE, and 2 distinct aspects per animal for EL, WM, and CTX. Differences between the three experiments were analyzed using one-way analysis of variance (ANOVA). To investigate the relationships between variables, simple linear regression and Pearson correlation analyses were performed (*p < 0.05)*.

## Acknowledgements

The authors would like to acknowledge Noah A Jasinkiewicz’s assistance with processing the MR images.

## Author contribution

R.T.B. conceived project, and contributed experimental design, surgical experiments, manuscript writing.

S.C.M. contributed to experimental design and surgical experiments.

R.R.M. performed animal husbandry and assisted with surgical experiments and experimental organization.

G.T.K contributed drug permeability and pharmacokinetic studies and analysis.

D.A. performed experiments related to immunofluorescence and confocal imaging, quantified imaging data, manuscript writing and figure construction.

F.O.B quantified MRI data, and constructed figures.

A.H. analyzed ICP data and contributed to manuscript writing and MRI data interpretation.

D.J.B. provided the TRPV4 antagonist and served as a liaison with GSK

A.D.C. Performed necropsy, tissue collection and histology

B,L.B-Y. conceived project, contributed to experimental design and manuscript writing provided and funding acquisition.

## Materials and Correspondence

All correspondence and material requests should be addressed to Bonnie Blazer-Yost.

Supplementary Information is available for this paper.

**Extended Table 1.** Mean ± SEM (mm^3^) ventricular volumes in Sham-operated pigs, control pigs induced with post-hemorrhagic hydrocephalus (PHH-C) and PHH pigs treated with GSK2789917 (PHH-Tx). Sham, n = 4. PHH-C, n = 7 (except n = 6 on Day 24). PHH-Tx, n = 6. The PHH-C pig with the largest ventricles pre-operatively (Day 1) was euthanized prior to the Day-24 MRI due to neurological deterioration.

|  | Day 1 | Day 10±1 | Day 17±1 | Day 24±2 |
|---|---|---|---|---|
| Lateral ventricles (excl. olfactory recesses) | Sham: 1408 ± 104.6<br>PHH-C: 1383 ± 124.5<br>PHH-Tx: 1350 ± 74.2 | Sham: 1549 ± 95.5<br>PHH-C: 4375 ± 541.7<br>PHH-Tx: 2683 ± 165.3 | Sham: 1812 ± 86.6<br>PHH-C: 4148 ± 495.3<br>PHH-Tx: 2672 ± 169.2 | Sham: 1911 ± 63.2<br>PHH-C: 3824 ± 365.3<br>PHH-Tx: 2547 ± 184.4 |
| Olfactory recesses | Sham: 945 ± 92.6<br>PHH-C: 1026 ± 84.9<br>PHH-Tx: 819 ± 63.3 | Sham: 977 ± 72.8<br>PHH-C: 1385 ± 93.7<br>PHH-Tx: 876 ± 77.7 | Sham: 972 ± 44.7<br>PHH-C: 1219 ± 98.5<br>PHH-Tx: 894 ± 103.0 | Sham: 981 ± 47.0<br>PHH-C: 972 ± 164.2<br>PHH-Tx: 784 ± 129.6 |
| Lateral ventricles (+ olfactory recesses) | Sham: 2353 ± 56.4<br>PHH-C: 2410 ± 212.1<br>PHH-Tx: 2169 ± 83.6 | Sham: 2527 ± 65.7<br>PHH-C: 5761 ± 583.6<br>PHH-Tx: 3559 ± 195.3 | Sham: 2783 ± 59.3<br>PHH-C: 5367 ± 471.8<br>PHH-Tx: 3566 ± 210.0 | Sham: 2891 ± 54.5<br>PHH-C: 4824 ± 360.1<br>PHH-Tx: 3330 ± 143.4 |
| 3<sup>rd</sup> ventricle | Sham: 295 ± 27.7<br>PHH-C): 254 ± 23.5<br>PHH-Tx: 249 ± 27.6 | Sham: 345 ± 23.5<br>PHH-C: 305 ± 45.3<br>PHH-Tx: 236 ± 20.0 | Sham: 383 ± 15.8<br>PHH-C: 353 ± 69.9<br>PHH-Tx: 231 ± 39.2 | Sham: 412 ± 17.6<br>PHH-C: 308 ± 35.0<br>PHH-Tx: 235 ± 20.8 |
| 4<sup>th</sup> ventricle | Sham: 998 ± 67.6<br>PHH-C: 863 ± 62.1<br>PHH-Tx: 851 ± 55.7 | Sham: 1088 ± 59.3<br>PHH-C: 1144 ± 147.2<br>PHH-Tx: 766 ± 57.0 | Sham: 1158 ± 59.4<br>PHH-C: 1304 ± 168.8<br>PHH-Tx: 664 ± 67.3 | Sham: 1220 ± 56.6<br>PHH-C: 1213 ± 104.5<br>PHH-Tx: 623 ± 70.6 |
| Total Ventricular Volume (all 4 ventricles) | Sham: 3645.5 ± 122.6<br>PHH-C: 3526.6 ± 267.8<br>PHH-Tx: 3269.9 ± 116.7 | Sham: 3959.3 ± 119.4<br>PHH-C: 7039.9 ± 691.0<br>PHH-Tx: 4561.6 ± 198.3 | Sham: 4324.0 ± 96.3<br>PHH-C: 7082.5 ± 498.7<br>PHH-Tx: 4460.8 ± 206.8 | Sham: 4514.3 ± 82.9<br>PHH-C: 6345.8 ± 353.6 (n=6)<br>PHH-Tx: 4188.9 ± 140.2 |

**Extended Data Figure 1.**
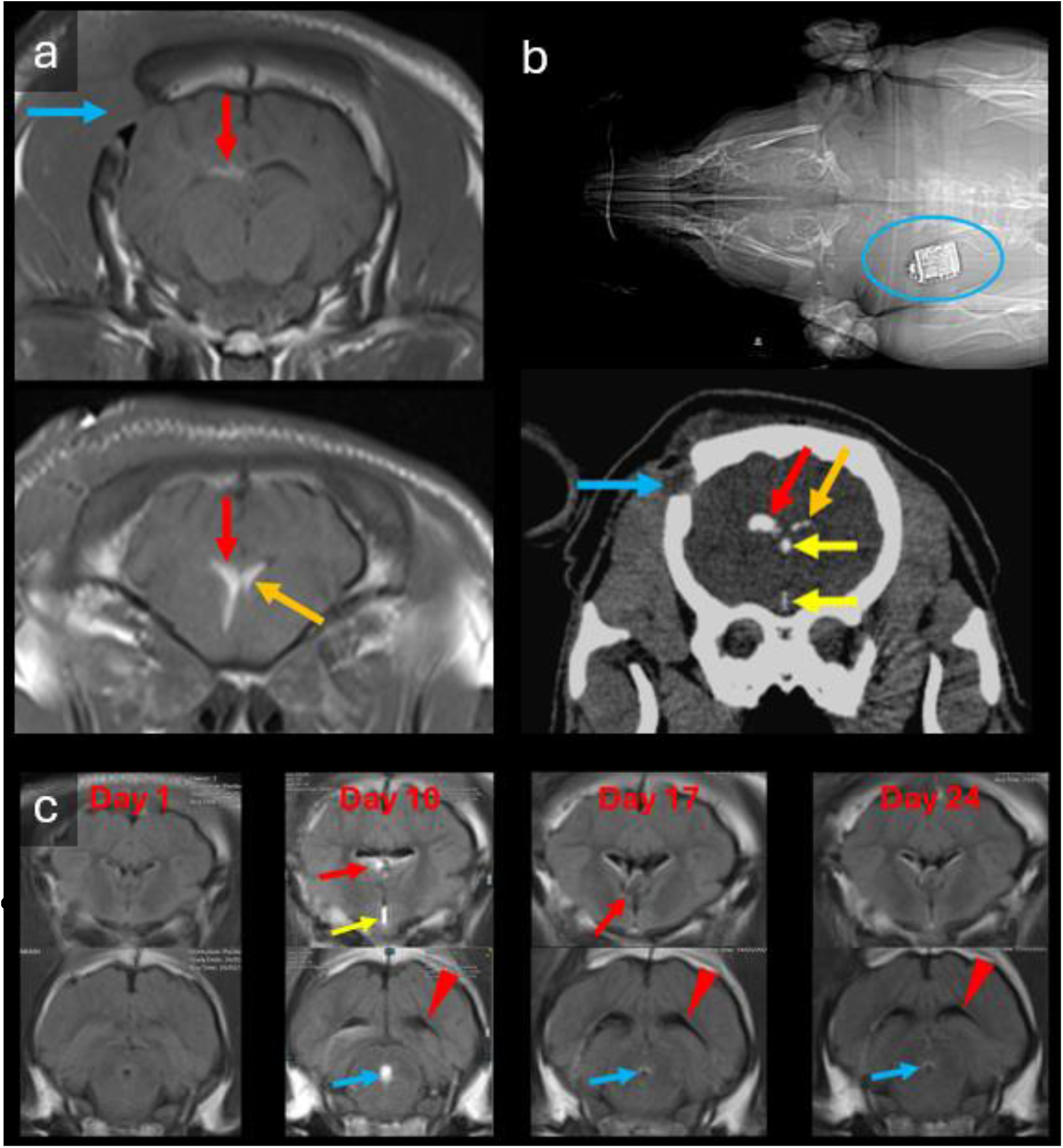
Figure 1a. Day-10 MRI confirming correct intraventricular injection of blood. T1-hyperintense (bright) blood has entered and enlarged the right ventricle (red arrows). Blood has entered the left ventricle (orange arrow). Blue arrow, rostral edge of the burrhole craniectomy. Transverse T1-weighted images at the level of the body (upper) and frontal horn (lower) of the lateral ventricle. Figure 1b. Day-2 CT confirming correct intraventricular injection of blood. Hyperattenuating (bright) blood has entered and enlarged the right ventricle (red arrow). Blood has also entered the left ventricle (orange arrow) and 3^rd^ ventricle (yellow arrows). Blue arrow, rostral edge of the burrhole craniectomy. The subcutaneous pressure sensor / battery pack unit (blue circle) is over the neck. Figure 1c. MRI on Day 1 (pre-op) to Day 24, of a PHH-Treatment pig induced with IVH and treated with GSK2789917 on Days 3 – 17. Transverse T2-weighted FLAIR images at the level of the interventricular foramina (top row) and the aqueduct (bottom row). Pre-operatively, the ventricles are small, containing only hypointense CSF. At Day 10, the lateral ventricles are enlarged, containing hyperintense hematoma (red arrows) and a smaller volume of hypointense CSF. The 3^rd^ ventricle is enlarged and contains hyperintense hematoma (yellow arrow). Some pigs displayed hyperintense material in the aqueduct (blue arrows). Ventriculomegaly persists at Days 17 and 24. At Day 17, ventricular and aqueductal hemorrhage is largely resolved; some hyperintensity persists in the right interventricular foramen (red arrow). Ventriculomegaly affects the posterior horns of the lateral ventricles on Days 10 – 24 (red arrow heads).

**Extended Data Figure 2.**
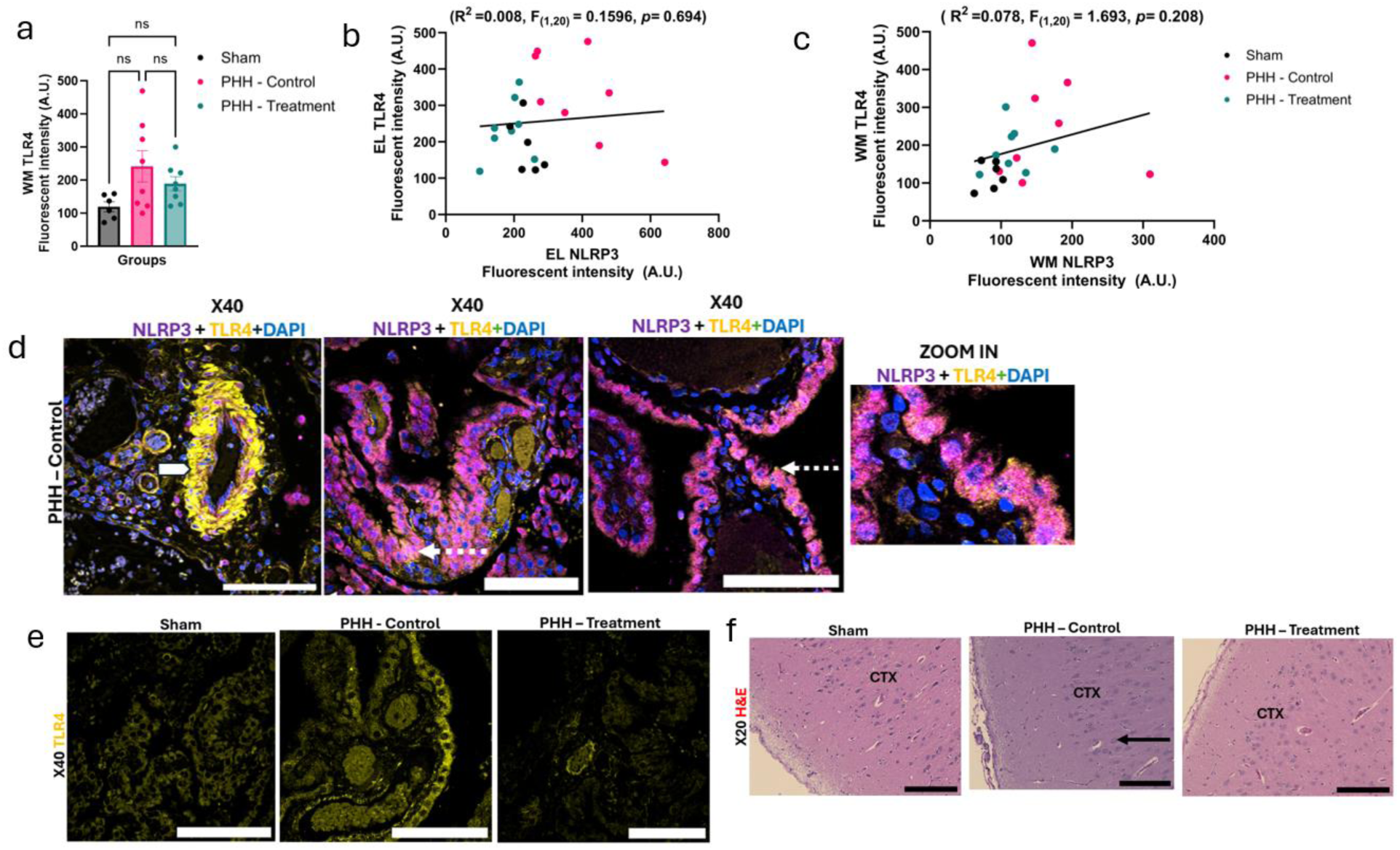
Regional analysis of TLR4 and NLRP3 expression in the ependymal lining, white matter, and cortex following PHH. (a) Quantification of white WM TLR4 fluorescence intensity (arbitrary units, A.U.). (b) Linear regression analysis of EL TLR4 fluorescence intensity *versus* EL NLRP3 fluorescence intensity (R² = 0.008, p = 0.694). (c) Linear regression analysis of WM TLR4 fluorescence intensity *versus* WM NLRP3 fluorescence intensity (R² = 0.078, p = 0.208). (d) Representative high magnification confocal immunofluorescence images of the choroid plexus tissue from PHH-Control animals stained for NLRP3 (red channel; displayed as magenta), TLR4 (green channel; displayed as yellow), and DAPI (blue). White block arrows indicate vascular TLR4 localization, while white dotted arrows indicate regions of NLRP3/TLR4 colocalization in the EL. Scale bar: 20 µm. (e) Representative confocal immunofluorescence images of TLR4 immunoreactivity (yellow) within the EL of Sham, PHH-Control, and PHH-Treatment animals. Scale bar: 20 µm. (f) Representative hematoxylin and eosin (H&E) images of the CTX in Sham, PHH-Control, and PHH-Treatment animals. The red arrow indicates enlarged cortical cells vacuolation in the PHH-Control group. Scale bar: 50 µm. Data are presented as mean ± SEM. **Abbreviations:** CTX, cerebral cortex; DAPI, 4′,6-diamidino-2-phenylindole; EL, ependymal lining; NLRP3, NOD-like receptor family pyrin domain-containing 3; PHH, post-hemorrhagic hydrocephalus; TLR4, Toll-like receptor 4; WM, white matter.

