## Supplemental Figures for "TRPV4 antagonist efficacy in a post-hemorrhagic hydrocephalus pig model"

Figure 1c. MRI on Day 1 (pre-op) to Day 24, of a PHH-Treatment pig induced with IVH and treated with GSK2789917 on Days 3 – 17. Transverse T2-weighted FLAIR images at the level of the interventricular foramina (top row) and the aqueduct (bottom row). Pre-operatively, the ventricles are small, containing only hypointense CSF. At Day 10, the lateral ventricles are enlarged, containing hyperintense hematoma (red arrows) and a smaller volume of hypointense CSF. The 3$^{rd}$ ventricle is enlarged and contains hyperintense hematoma (yellow arrow). Some pigs displayed hyperintense material in the aqueduct (blue arrows). Ventriculomegaly persists at Days 17 and 24. At Day 17, ventricular and aqueductal hemorrhage is largely resolved; some hyperintensity persists in the right interventricular foramen (red arrow). Ventriculomegaly affects the posterior horns of the lateral ventricles on Days 10 – 24 (red arrow heads).

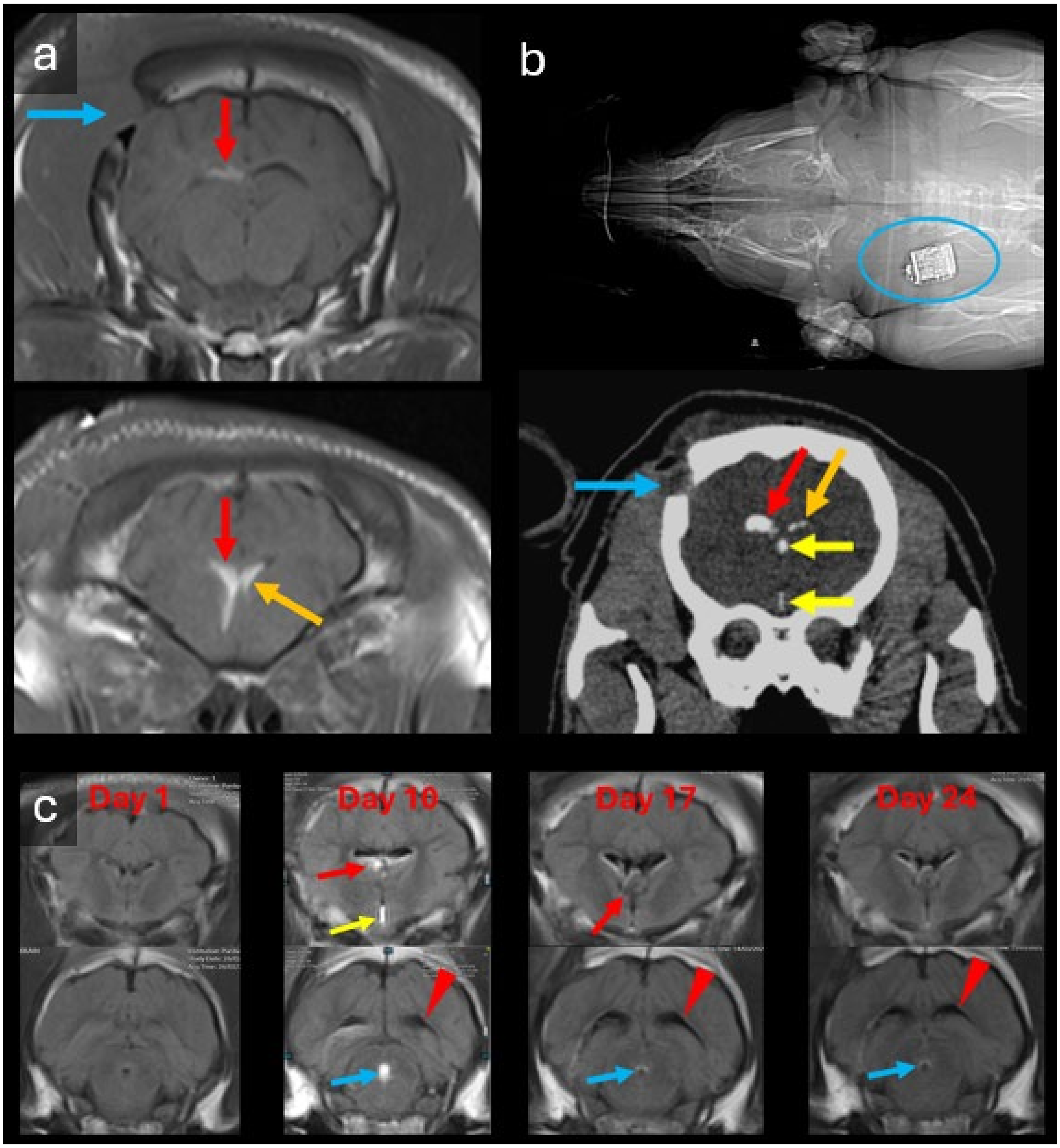
a
b
c
Day 1
Day 10
Day 17
Day 24

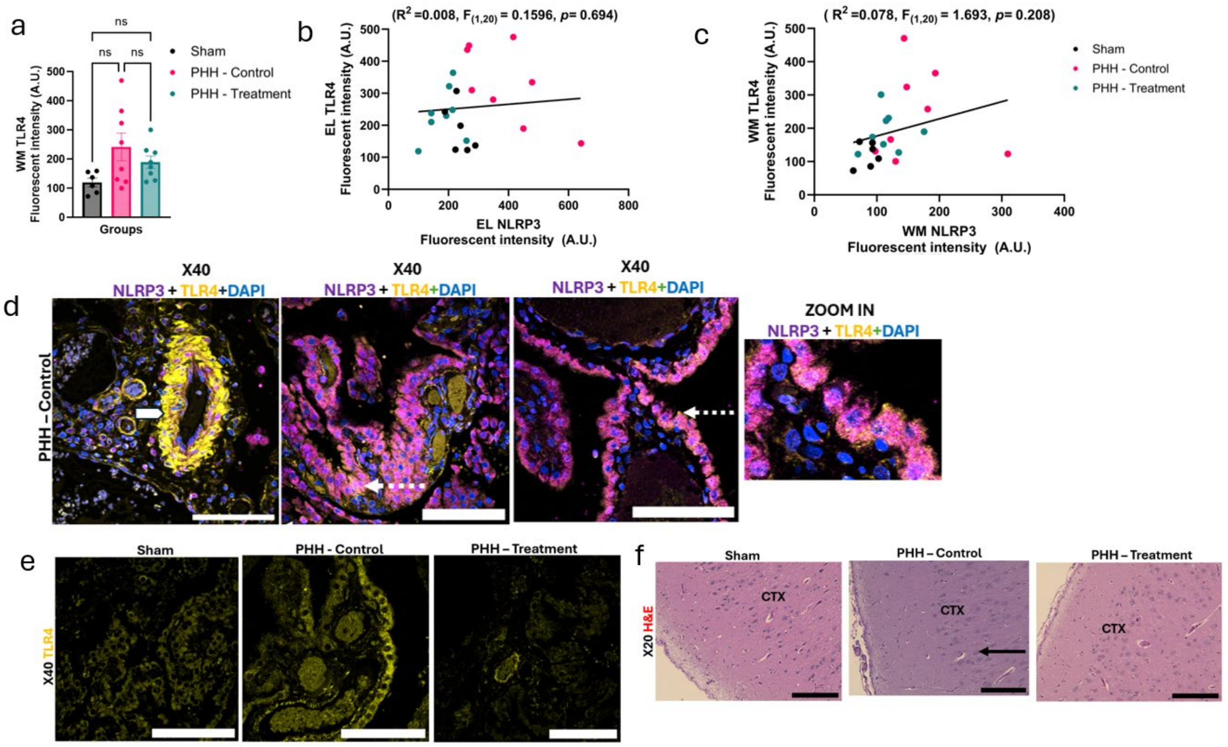

**Extended Data Figure 2. Regional analysis of TLR4 and NLRP3 expression in the ependymal lining, white matter, and cortex following PHH.**

(a) Quantification of white WM TLR4 fluorescence intensity (arbitrary units, A.U.).

(b) Linear regression analysis of EL TLR4 fluorescence intensity *versus* EL NLRP3 fluorescence intensity ($R^2$ = 0.008, p = 0.694).

(c) Linear regression analysis of WM TLR4 fluorescence intensity *versus* WM NLRP3 fluorescence intensity ($R^2$ = 0.078, p = 0.208).

(d) Representative high magnification confocal immunofluorescence images of the choroid plexus tissue from PHH-Control animals stained for NLRP3 (red channel; displayed as magenta), TLR4 (green channel; displayed as yellow), and DAPI (blue). White block arrows indicate vascular TLR4 localization, while white dotted arrows indicate regions of NLRP3/TLR4 colocalization in the EL. Scale bar: 20 µm.

(e) Representative confocal immunofluorescence images of TLR4 immunoreactivity (yellow) within the EL of Sham, PHH-Control, and PHH-Treatment animals. Scale bar: 20 µm.

(f) Representative hematoxylin and eosin (H&E) images of the CTX in Sham, PHH-Control, and PHH-Treatment animals. The red arrow indicates enlarged cortical cells vacuolation in the PHH-Control group. Scale bar: 50 μm.

Data are presented as mean ± SEM.

**Abbreviations:** CTX, cerebral cortex; DAPI, 4',6-diamidino-2-phenylindole; EL, ependymal lining; NLRP3, NOD-like receptor family pyrin domain-containing 3; PHH, post-hemorrhagic hydrocephalus; TLR4, Toll-like receptor 4; WM, white matter.
